# Dynamic decoding of VEGF signaling and coordinated control of multiple phenotypes by the Src-TEM4-YAP pathway

**DOI:** 10.1101/2023.10.19.562360

**Authors:** Sung Hoon Lee, Tae-Yun Kang, Andre Levchenko

## Abstract

Responses of endothelial cells to elevated levels of Vascular Endothelial Growth Factor (VEGF), frequently accompanying local decrease in oxygen supply, include loosening of cell contacts, rearrangement of cells in the process of vessel remodeling and ultimately, angiogenic growth. How these complex processes, occurring on diverse time scales, are coordinated and how they are guided by a single key signaling input, is still incompletely understood. Here we show that the various phenotypic responses associated with VEGF signaling are controlled at different steps of a pathway involving sequential activation of Src, TEM4, YAP and components of pro-angiogenic Notch signaling. Strikingly, due to feedback regulation at different pathway levels, the functional outcomes are controlled by oscillations of the pathway components occurring on distinct time scales. Deeper pathway layers integrate faster upstream responses and control progressively slower phenotypic outcomes. This signal decoding pathway organization can ensure a high degree of complexity in a vital physiological process.

## Introduction

Multiple developmental and physiological processes, such as angiogenesis, i.e., formation of new blood vessels from the existing ones, involve complex tissue reorganization and coordinated cell responses to a handful of signaling cues. In angiogenic sprouting, these responses include regulation of cell contacts and endothelia barrier, local cell re-arrangement, cell fate specification, cell migration and proliferation. Furthermore, genetic, biochemical and cell biological evidence have pointed to a single molecular factor, vascular endothelial growth factor (VEGF), as the key input that guides and orchestrates all these processes (although other factors play important additional modulating roles). Understanding how the signaling networks triggered by a single ligand can regulate diverse but coordinated phenotypic responses, occurring on multiple temporal scales is a key challenge of quantitative and systems biology analysis of signal transduction.

Loosening of cell contacts is a key feature of and a prerequisite for distinct responses to VEGF and other angiogenic stimuli. In the short term, it can facilitate an increased exchange of the chemical and cellular components between the bloodstream and the local hypoxic and possibly inflamed tissue. In the longer term, it can underlie reorganization of the endothelium architecture, enabling alterations of mutual cell positions prior and during the initiation of angiogenic sprouting. Animal and other studies have strongly suggested that the key VEGF-activated signaling molecule controlling the stability of cellular junctions in the endothelium is Src (1). However, how VEGF-Src signaling can influence diverse phenotypic responses occurring on diverse time scales is largely uncharacterized. Even the analysis of how VEGF controls the endothelial barrier function has been plagued by controversies and open questions (2). In particular, although it has been originally assumed that phosphorylation of VE-cadherin by Src is responsible for this effect of VEGF, it has been later shown that this phosphorylation event is not sufficient to explain the increased vascular permeability (3). Furthermore, it is not clear whether VEGF-stimulated Src activity is transient and only provides a brief trigger of the longer term angiogenic effects dependent on loosening cellular contacts, or if persistent Src activity is required for VEGF responses occurring over multiple days.

In this study, we examined the dynamics of Src activation, its decoding by the downstream signaling network and its influence on diverse phenotypic responses in the monolayer endothelium model. Although this model is a simplistic representation of complex in vivo endothelium, it permits a careful examination of the biochemical and genetic interactions in the VEGF-Src activated network. We find that Src activity is oscillatory, with this oscillating signal decoded by different layers of the signaling networks on different time scale. These results provide an integrated picture of how a single dynamic signaling input can be converted into complex dynamic outputs on different times scales controlling multiple phenotypic outcomes.

## Results

### VEGF triggers oscillatory Src activation and dynamic TEM4 expression controlling the barrier function in endothelial monolayers

To explore the pSrc dependent control of endothelial phenotypes (Fig. 1A), we experimented with the monolayer of human umbilical vein endothelial cells (HUVEC). Although, the endothelial monolayer models clearly cannot reproduce the complexity of *in vivo* blood vessels, they have been successfully used in the past to capture some of the key aspects of signaling and regulatory s stimulinteractions in mature and developing endothelia (4). We found that, in response to continuouation with 100 ng/ml VEGF, phosphorylated and total Src levels underwent oscillatory dynamics, with the ratio of phosphorylated to total Src (pSrc/Src) also displaying persistent oscillations with approximately 3.5 hr. period (Fig. 1B). These oscillations (also observed with reduced amplitude for a lower VEGF stimulation dose (Fig. S1A)), were correlated with an oscillatory phosphorylation of VE-cadherin in the same cells, consistent with Src mediated phosphorylation of this protein. The VE-cadherin phosphorylation is assumed to be critical for loosening of the endothelial adherens cell junctions and increased monolayer permeability (1). Interestingly, we also observed periodic increases in cell motility, leading to re-arrangement of cell positions with respect to each other in response to VEGF (Fig. 1C). These re-arrangements provided indirect support to the overall loosening of cell contacts, leading to an increased ‘fluidity’ of the monolayer. Importantly, the peaks of migration speed coincided with the peaks of Src phosphorylation. Inhibition of Src activity by Dasatanib led to both inhibition of cell motility speed and abrogation of its oscillatory nature, further supporting the critical role of Src in the process of VEGF induced cell rearrangement. Of note, the prominent effectors of VEGF, Erk and Akt, did not exhibit oscillatory patterns over time (Fig. S1B). Further, both PI3K and MEK inhibitors do not interfere with the oscillatory changes in cell migration speed, although both inhibitors negatively impact the speed magnitude during treatment (Fig. S1C). This data implies that VEGF induces oscillatory Src activity, playing a crucial role in orchestrating the temporal coordination of the oscillatory migration patterns.

**Figure 1.**
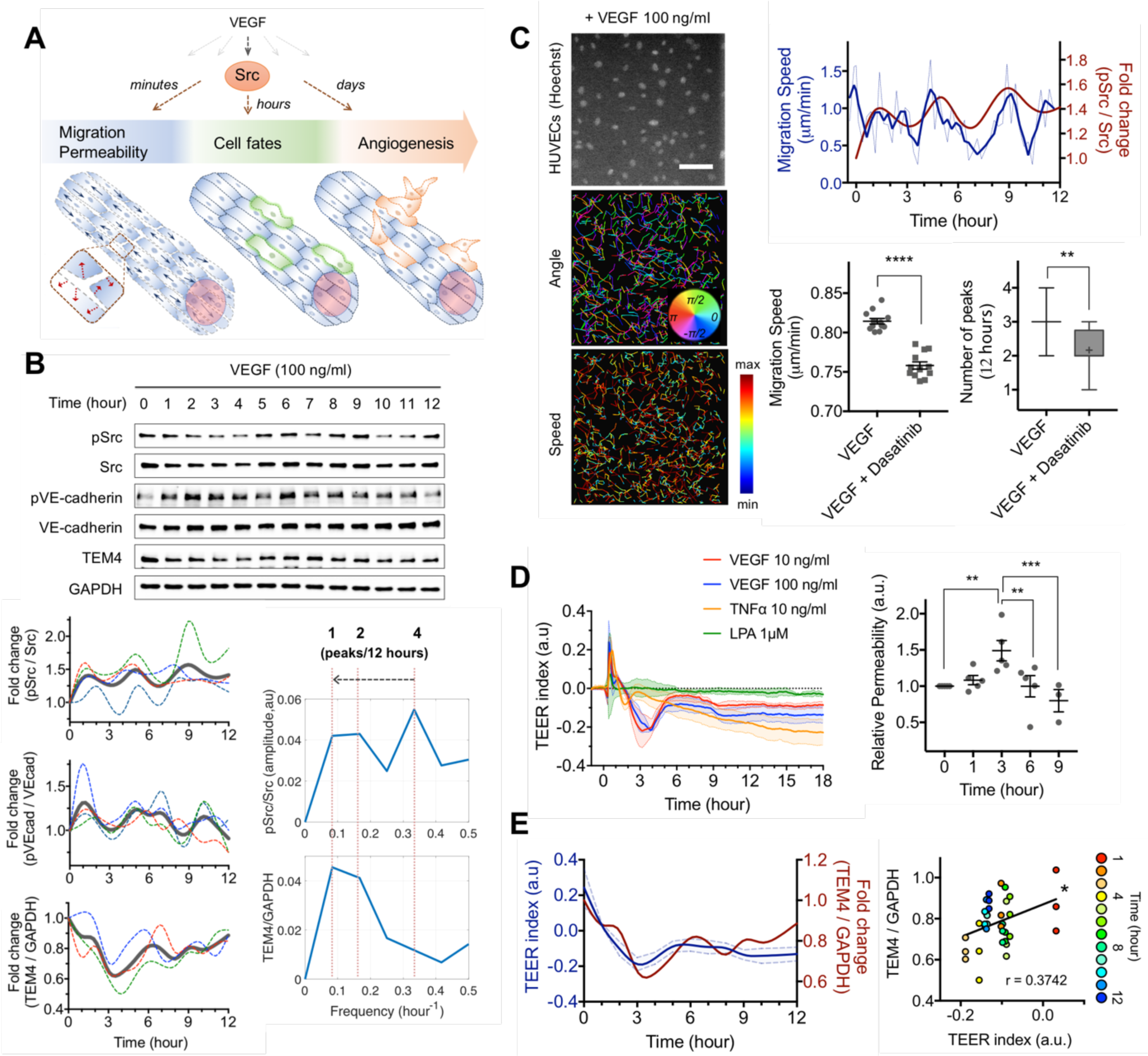
VEGF controls multiple phenotypic processes, with each process possessing a distinct signaling frequency. **(A)** Schematic of a process controlling multiple biological activities including cell migration, junction permeability, cell fates and angiogenesis, mediated by VEGF. **(B) (top)** Representative immuno-blot results of pSrc (Tyr416), Src, pVE-cadherin (Tyr685), VE-cadherin and TEM4 of HUVECs. GAPDH was used as a loading control. **(bottom, left)** Fold change analysis of the results (n=4 (pSrc/Src, pVEcad/VEcad), n=3 (TEM4/GAPDH)). Each color dotted line represents individual experiments and gray lines indicate the average. **(bottom, right)** Fast Fourier Transform analysis based on the time domain results in the middle column. **(C) (left)** A representative fluorescent image of HUVECs dyed with Hoechst to visualize nucleus, and results of an automated tracking analysis representing angle and speed of trajectories of Hoeschst from images taken every 15 minutes for 12 hours. **(right, top)** Migration speed of HUVECs in response to VEGF 100 ng/ml (blue). The thin lines represent averaged migration speed from individual cells and the thick lines indicate moving average of the thin lines. Temporal kinetics of pSrc/Src from Figure 1B is overlaid (red). **(right, bottom left)** Average migration speed for 12 hours of the cells in response to the 30-min pretreatment by Dasatinib 10 nM, LY294002 1 μM (PI3K inhibitor), or PD0325901 1 nM (MEK inhibitor), followed by the treatment of VEGF 100 ng/ml. (****P<0.0001, n=12 field of views (FOVs) for all conditions, n=447 cells tracked in average for control, n=521 cells tracked in average for Dasatinib, n=433 cells tracked in average for LY294002, n=405 cells tracked in average for PD0325901, all time points included.) **(right, bottom right)** The number of peaks detected using the peak detection analysis for each condition. The data covers min to max, with + indicating the mean value. (**P=0.0015) The following threshold values were used for the analysis, AmpT=0.5, SlopeT=0.000099, SmoothW=5 and FitW=4) **(D) (left)** Relative real-time impedances of mono-layered HUVECs in response to VEGF 10, 100 ng/ml, TNFα 10 ng/ml and LPA 1 μM. Each data was subtracted from control impedance (no treatment, serum-starved) representing baseline 0, with error bar representing s.e.m. (n=3). Rapid increase of impedances in VEGF condition within 30 minutes is due to an increase of cell area, as confirmed in **Figure S1D**. **(right)** FITC-dextran results. Fluorescent intensity measurement over time from HUVECs in response to VEGF 100ng/ml. Data were analyzed by two-way ANOVA with error bar representing s.e.m. (0 vs. 3 hours (**P=0.0092), 3 vs. 6 hours (**P=0.0089) and 3 vs. 9 hours (***P=0.0009), n=5 (0,1,3,6 hours), n=3 (9 hours)) **(E) (left)** Immuno-blot results of TEM4 (blue) from (Figure 1B) overlaid in the impedance measurement results (red). A starting point at time 0 for TEER index was adjusted to when the index is the highest. **(right)** Scatter plot of fold changes of TEM4/GAPDH from (Figure 1B) over TEER index measured in (Figure 1D) (*P=0.0246). Color indicates time exposed to VEGF 100 ng/ml.

To evaluate the monolayer permeability more directly, we measured the Trans-Epithelial Electrical Resistance (TEER) (5). We found that the permeability indeed showed a dynamic increase following VEGF application. However, strikingly, the dynamics of this permeability change were distinct from the oscillatory pSrc or VE-cadherin dynamics. In particular, instead of 3-4 peaks over 12 hrs., we found a single pronounced dip in the monolayer impedance, followed by partial recovery (Fig. 1D, S1E). We then investigated another protein implicated in controlling endothelial permeability, a cell junction-associated Rho guanine exchange factor (GEF), tumor endothelial marker 4 (TEM4) (6). We found that the abundance of this protein in response to VEGF stimulation was also dynamically altered, and that this dynamics had a striking similarity to the dynamics of TEER (Fig. 1E, S1F). This result suggested that it was TEM4 that largely defined the endothelial permeability in this experimental model. This conclusion was supported by multiple other pieces of evidence. First, we found no VEGF-induced changes in Src association with other Rho-GEF or -GAP proteins implicated in regulation of adherens or tight junctions, including GEF-H1, p114 RhoGEF, PDZ RhoGEF, Rich1 and p190 RhoGAP (6–13), suggesting that Src-mediated effects in RhoA activity are TEM4 specific (Fig. S2A). Subsequently, we analyzed GEF-H1 in various experiments as a control, since inhibition of GEF-H1 is reported to be particularly important for stabilizing junction organization (7). In particular, we further found that up-regulation of both RhoA activity and activation of the RhoA effector mDia1 were TEM4-specific, but independent of GEF-H1 (Fig. S2B,C). Furthermore, TEM4, but not GEF-H1, was upregulated at the plasma membrane at high cell densities (Fig. S2D,E). Also, as expected, TEM4 was associated with and was stabilized by β-catenin, further pointing to the junctional localization (Fig. S2F) (6, 12, 13). Most importantly, interference with TEM4 expression led to a much greater and dynamically persistent increase of the monolayer permeability vs. the control case (Fig. S2G). Overall, these data suggested that TEM4 is a key dynamic determinant of the monolayer permeability following VEGF stimulation.

These findings raised several questions. First, what is the mechanism controlling the oscillatory Src dynamics? Second, does the TEM4 dynamics depend on Src, in spite of having a distinct temporal regulation profile? And if so, what mechanism controls the overall TEM4 dynamics? We addressed these questions in the following experimental analysis.

### Src activity oscillations are controlled by negative feedback involving Csk

A common cause of oscillatory dynamical responses is the presence of negative feedback interactions (14). We thus focused on a key negative Src regulator, inhibiting Src phosphorylation on Tyr416, the C-terminal Src kinase (Csk) that has been suggested as a putative negative feedback mediator (15). We transfected the cells with a constitutively active Src mutant (caSrc) and measured the expression level of Csk. In addition, we abrogated Csk expression by siRNA and determined the basal level of Src activity. We found that an increased Src activity upregulated Csk expression, and Csk down-regulated Src activity (Fig. 2A), providing experimental evidence for the Src-Csk negative feedback regulation. A simple ordinary differential equations (ODE) model suggested that,if the Csk-mediated signaling feedback indeed controls the Src dynamics, Csk concentration is also expected to oscillate with the same frequency but a phase shift vs. Src phsophorylation (Fig. 2B,C). To explore this dynamic effect, we immuno-blotted the levels of Csk along with phosphorylated Src (Tyr416) and compared them in VEGF-stimulated cells. We found that Csk level indeed oscillated in response to VEGF, with the same frequency and with a time delay vs Src phsophorylation dynamics (Fig. 2D). These results, in combination, suggested that the likely mechanism of oscillatory Src activity dynamics is its negative feedback interaction with Csk.

**Figure 2.**
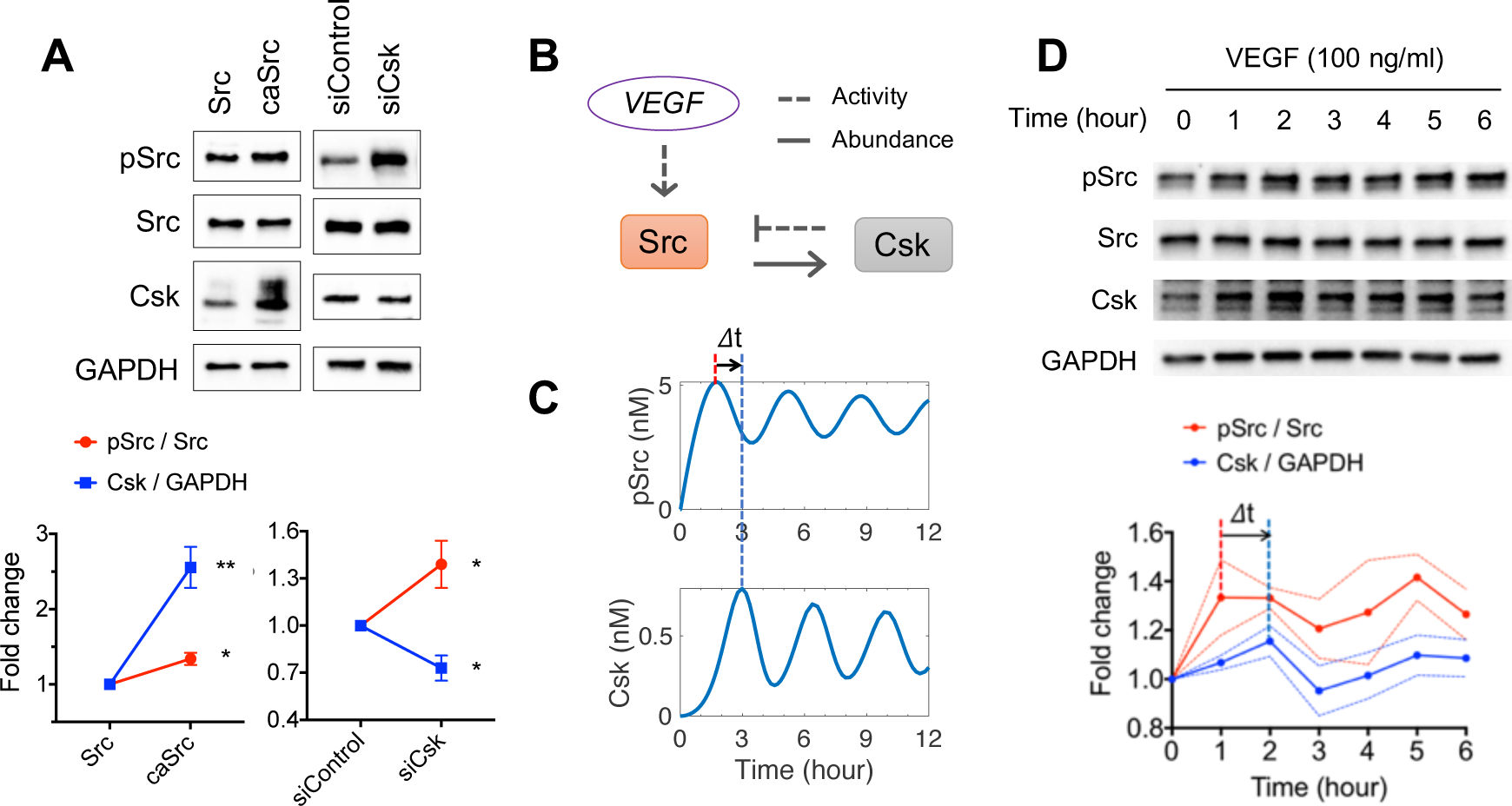
A negative feedback between pSrc and Csk leads to oscillatory dynamics. **(A)** Representative immuno-blot results of pSrc (Tyr416), Src and Csk of HUVECs infected with Src overexpression (Src), a mutation that is constitutively active (caSrc), control siRNA and Csk siRNA, and fold change analysis. GAPDH was used as a loading control. (For Src vs. caSrc, pSrc/Src *P=0.0132 (n=3), Csk/GAPDH **P=0.0012 (n=4), for siControl vs. siCsk, pSrc/Src *P=0.032 (n=3), Csk/GAPDH *P=0.0279 (n=3)) **(B)** Schematic diagram summarizing VEGF induced pSrc with Csk. **(C)** Model prediction of temporal changes of pSrc, and of Csk. **(D)** Representative immuno-blot results of pSrc (Tyr416), Src and Csk of HUVECs in response to VEGF 100 ng/ml, and fold change analysis. (n=4)

### Src activity leads to down-regulation of TEM4 levels

We next addressed whether TEM4 expression dynamics is Src dependent and thus could be responsive to oscillatory Src dynamics. Hypothetically, VEGF can regulate TEM4 through a range of mechanisms that influence the junctional integrity, which may or may not include Src. In particular, in addition to to Src, VEGF can relay the signals via Vav2, Rac and Pak, resulting in endocytosis of phosphorylated VE-cadherin (1). Also, VEGF promotes FAK activation, leading a localization of FAK to VE-cadherin and ultimately junction disassociation through phosphorylation of β-catenin at Y142 (16). To explore which, if any, of these mechanisms may be involved, we transfected HEK 293T cells with a GFP-TEM4 encoding plasmid and co-immunoprecipitated the GFP-TEM4 fusion protein with Src, Vav2, Pak and FAK. Compared to GFP-empty expressing cells, only Src was immuno-precipitated with GFP-TEM4 (Fig. 3A). Furthermore, consistent with these co-immunoprecipitation (IP) results, immunostaining suggested that Src was co-localized with TEM4 at cell-cell junctions, suggesting that Src indeed forms a complex with TEM4 at these locations (Fig. 3B).

**Figure 3.**
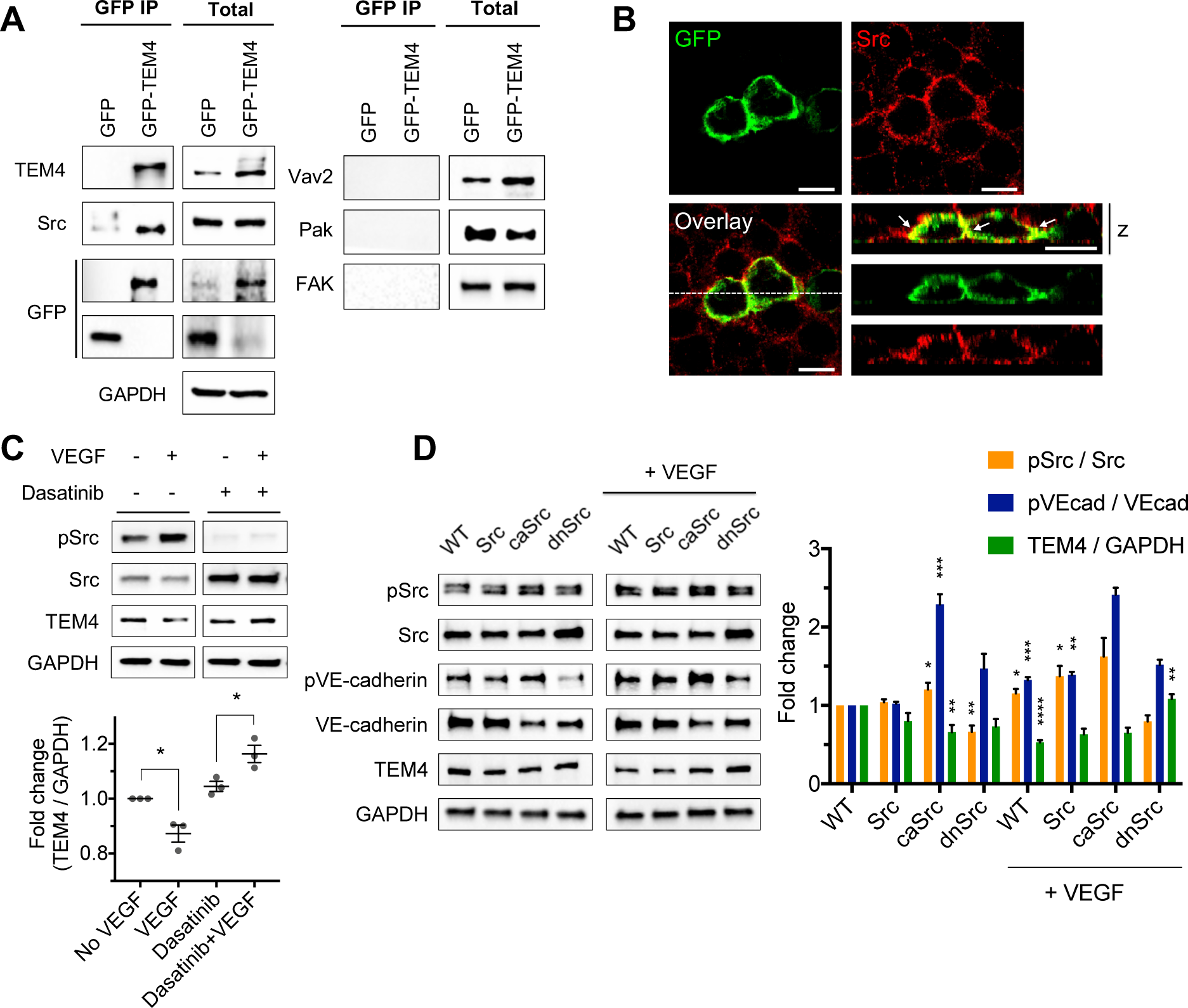
TEM4 down-regulation is mediated by pSrc activity. **(A)** Representative immuno-blot results of co-immunoprecipitation of GFP against TEM4 and Src, Vav2, Pak and FAK. GAPDH was used as a loading control. (n=3) **(B)** Representative immuno-staining results of Src (red) in mono-layerd TEM4-GFP (green) expressing HEK 293T cells. Cross section (z) view of cells from white dotted lines. White arrows indicate co-localization of Src with TEM4 (scale bar: 10 μm). **(C)** Representative immuno-blot results of pSrc (Tyr416), Src and TEM4 in response to the 15-hour pretreatment of DMSO or Dasatinib 10 nM, followed by the treatment of VEGF 100 ng/ml for 2 hours, and fold change analysis. GAPDH was used as a loading control. (For No VEGF vs. VEGF, *P=0.0147 (n=3), for Dasatinib vs. Dasatinib+VEGF, *P=0.0314 (n=3)) **(D)** Representative immuno-blot results of pSrc (Tyr416), Src, pVE-cadherin (Tyr685), VE-cadherin and TEM4 in response to VEGF 100 ng/ml (2 hours) for HUVECs of wild-type (WT), Src overexpression (Src), a constitutively active Src mutant (caSrc) and a dominantly negative Src mutant (dnSrc), and fold change analysis. GAPDH was used as a loading control. P-values for graph are as follows: caSrc (pSrc/Src *P=0.0429, pVE-cadherin/VE-cadherin ***P=0.0006, TEM4/GAPDH **P=0.0029), dnSrc (pSrc/Src **P=0.0029), WT with VEGF (pSrc/Src *P=0.0295, pVE-cadherin/VE-cadherin ***P=0.00082, TEM4/GAPDH ****P<0.0001), Src with VEGF (pSrc/Src *P=0.038, pVE-cadherin/VE-cadherin **P=0.00139) and dnSrc with VEGF (TEM4/GAPDH **P=0.0015) with the following number of experiments, pSrc/Src (n=5), pVE-cadherin/VE-cadherin (n=3) and TEM4/GAPDH (n=7).

Next, we tested whether Src activation leads to a change in TEM4 abundance. We applied 10 nM of a Src inhibitor, Dasatinib, to fully confluent HUVECs for 2 hours. This treatment completely rescued VEGF-mediated TEM4 down-regulation (Fig. 3C). We complemented this pharmacological perturbation with genetic ones, by exploring cells retrovirally infected with Src mutants including constitutive active Y527F (caSrc), dominant negative K295R (dnSrc) or wild type control (Src). As expected, Tyr416-phosphorylated Src normalized to total Src was highly elevated in caSrc cells compared to wild-type controls, but was down-regulated in dnSrc cells. Further, we found that only caSrc expressing cells displayed a significantly lower TEM4 expression levels, even without VEGF treatment. On the other hand, even following VEGF treatment for 2 hours, there was no TEM4 down-regulation in dnSrc cells (Fig. 3D). Consistently with this finding, VEGF-treatment in dnSrc HUVECs did not result in an increased junctional permeability. Furthermore, we found that treatment of cells with TNF, another ligand loosening endothelial junctions, resulted in lack of Src activating and TEM4 down-regulation (Fig. S3A,B). Taken together, this data indicates that Src activity is indeed directly involved in VEGF-mediated TEM4 down-regulation.

### The recovery of junction permeability is controlled by a negative feedback loop involving TEM4-regulated YAP activity and YAP-driven *TEM4* transcription

Different dynamic profiles of Src phosphorylation vs TEM4 degradation and re-synthesis suggest that TEM4 abundance does not simply kinetically follow Src activity, but is affected by an independent and slower process. Because TEM4 is a transcriptional target of YAP-TEAD in breast cancer cell lines (MDA-MB-231) (17), it is possible that the TEM4 re-synthesis (following pSrc-mediated, VEGF-induced degradation) and the ensuing junctional recovery are also mediated through a transcriptional regulation by YAP in endothelial cells. Moreover, this regulation might occur in a feedback fashion, because TEM4 is required for stable junction formation and barrier integrity ((6) and our data above) and YAP activity is directly affected by the degree of junction formation or changes of actin organization (Fig. S4A) (18–20). Therefore, YAP may control TEM4 on the transcriptional level, whereas TEM4 may in turn regulate YAP activity through junction stabilization. To test this hypothesis, we lentivirally knocked down TEM4 and compared intracellular YAP localization at different cell densities by immunostaining. We again used similar genetic GEF-H1 manipulation as a control. Our results indicate that YAP is mostly localized in cell nuclei at a low or intermediate cell densities, but is both nuclear and cytosolic at high cell density, in both control and GEF-H1 knockdown cells. However, YAP displayed nuclear localization in both high and lower cell density cultures in TEM4 knockdown cells, indicative of high activity due to junctional destabilization in confluent cells (Fig. 4A). We further found that this increased nuclear YAP translocalization in TEM4 knockdown cells was correlated with an increased cell area and FAK activity (Fig. 4B,C) (21, 22), which are both expected to contribute to YAP activation under the conditions of loosened cell contacts.

**Figure 4.**
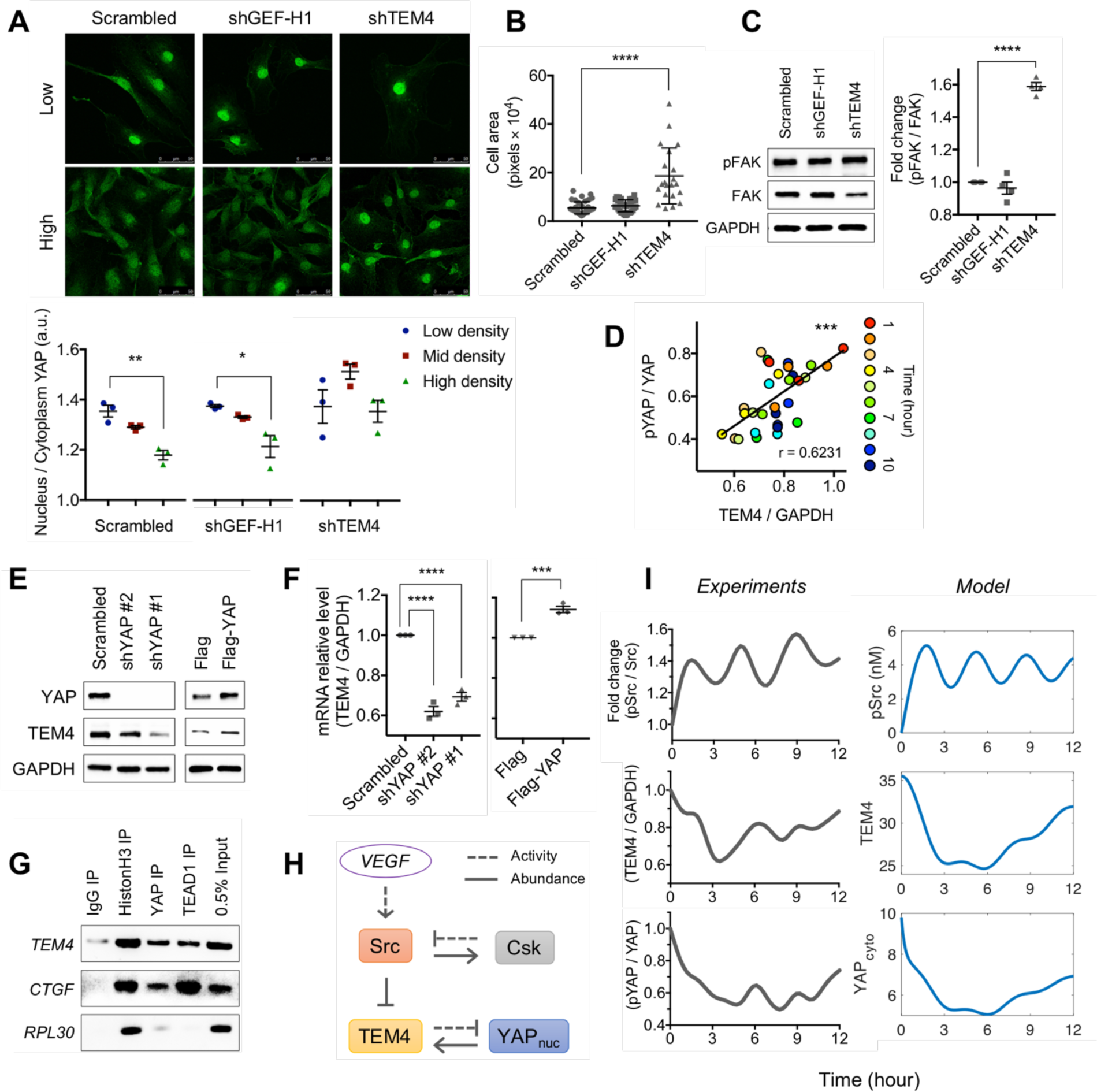
TEM4 down-regulation leads to YAP trans-localization into nucleus, leading to *TEM4* transcription. **(A) (top)** Representative immuno-staining results of YAP for low and high densities of HUVECs infected with Scrambled shRNA, shGEF-H1 and shTEM4 (scale bar: 50 μm). **(bottom)** Quantified localization of YAP nucleus over cytoplasm for low, mid and high density of HUVECs infected with Scrambled shRNA, shGEF-H1 and shTEM4. (Scrambled (**P=0.004355), shGEF-H1 (*P=0.02173), n=3) **(B)** Cell area of HUVECs infected with Scrambled shRNA (n=40), shGEF-H1 (n=31), shTEM4 (n=21). (****P<0.0001) **(C)** Representative immuno-blot results of pFAK (Tyr397) and FAK of HUVECs infected with Scrambled shRNA, shGEF-H1 and shTEM4, and fold change analysis. GAPDH was used as a loading control. (****P<0.0001, n=4) **(D)** Scatter plot of fold changes of TEM4/GAPDH over pYAP/YAP, based on immuno-blot results including Figure 1B and **Figure S4D** (***P=0.0002). Color indicates time exposed to VEGF 100 ng/ml. **(E)** Representative immuno-blot results of YAP and TEM4 of HUVECs infected with Scrambled shRNA, shYAP#2 and shYAP#1 and of HUVECs transfected with Flag-empty, YAP-Flag. GAPDH was used as a loading control. (n=3) **(F)** mRNA relative level of TEM4 of HUVECs infected with Scrambled shRNA, shYAP#2, shYAP#1 and of HUVECs transfected with Flag-empty and YAP-Flag. (***P=0.0009, ****P<0.0001, n=3) **(G)** Representative PCR results of chromatin-immunoprecipitation of *TEM4*, *CTGF* and *RPL30* against IgG, HistonH3, YAP, TEAD1 with 0.5% input lysates. (n=3) **(H)** Schematic diagram showing VEGF induced pathway involving Src and TEM4-YAP. **(I)** Experimental results (**left**, based on immuno-blot results including Figure 1B and **Figure S4D**) and model prediction **(right)** of pSrc, TEM4 and cytoplasmic YAP.

According to our hypothetical mechanism, the feedback-based re-synthesis of TEM4 does not initiate till after 2 hrs of VEGF stimulation, with the initial kinetics of TEM4 (and therefore YAP activity) driven entirely by the Src input over this initial time period. We thus expected both TEM4 and pYAP (the Ser127-phosphorylated, inactive, cytoslic and junction-localized form of YAP) to decrease over the first 2 hrs of VEGF input. We indeed observed this decrease, with the levels of pYAP and TEM4 being highly correlated over this time and subsequent period of VEGF stimulation (Fig. 4D, S4B). Furthermore, since the amplitude of pSrc and thus TEM4 oscillations were VEGF dose dependent, we expected the same dose dependency to be valid for pYAP, which we indeed observed over a few orders of VEGF magnitude, at 2 hrs of VEGF stimulation (Fig S4C,D). Overall, these results support the hypothesis of YAP activation due to TEM4 degradation during the early stages of VEGF input.

Our hypothesis also postulates delayed transcriptionally controlled YAP-mediated TEM4 re-synthesis, which thus closes the negative TEM4-YAP feedback loop during the later stages of 12 hr. VEGF stimulation. To investigate this part of the mechanism, we used LPA as a YAP inducer (20), because LPA does not disrupt cell-cell junctions or cause TEM4 down-regulation (Fig. 1D). Indeed, pYAP levels were down-regulated following LPA treatment, indicating an increase in the transcriptionally active YAP (Fig. S5A). Importantly, the treatment by LPA also led to an increase of TEM4 protein level (Fig. S5B), supporting our hypothesis. We then inhibited the transcriptional activity of YAP, using Verteporfin, a drug that suppresses YAP-TEAD1 interaction. We found that this treatment abrogated TEM4 synthesis following LPA treatment as well as basally, suggesting that inhibition of YAP-TEAD1 binding affects TEM4 up-regulation (Fig. S5C). Consistently, Verteporfin (+ VEGF) treatment also led to a persistent increase in the monolayer permeability over 12 hrs., without any indication of a partial recovery of the barrier function (Fig. S5D), suggesting that YAP transcriptional activity plays a key role in mediating the recovery of permeabilized cell-cell junctions.

We sought to support these findings by genetic perturbations of YAP abundance by using YAP shRNA mediated knockdown and YAP overexpression. We found that both mRNA and protein levels of TEM4 were significantly downregulated following YAP knockdown in HUVECs and HEK293T cells, whereas overexpression of YAP led to significant up-regulation of TEM4 mRNA and protein levels (Fig. 4E,F). Furthermore, chromatin-immunoprecipitation further revealed that both YAP and TEAD1 were bound to the promotor region of *TEM4*, supporting direct transcriptional regulation by YAP-TEAD1 complexes (Fig. 4G).

These results provided evidence for our hypothesis regarding TEM4 regulation, suggesting that it is controlled by pSrc input and a feedback regulation by YAP. The transcriptional nature of the feedback makes it relatively slower than the dynamics of pSrc regulation based on Src-Csk feedback. To describe how both Src and YAP activities can combine in defining TEM4 and, thus, monolayer permeability dynamics, we extended the ODE model to include the TEM4-YAP feedback downstream of Src (Fig. 4H,I). We trained the model using parameters taken from the previous literatures (23–25), or obtained through the data-fitting from experimental dynamics of pSrc, TEM4 and pYAP. The close agreement between the model and experimental findings supported the feasibility of the proposed mechanism. We next tested the model to explore cell responses that the model was not ‘trained’ to describe — focusing on studying the effects of transient VEGF stimulation (Fig. S5F). This analysis was particularly instructive for addressing the question of the importance of multiple peaks in Src phosphorylation in controlling the complex downstream biochemical and phenotypic responses. We thus first simulated and then experimentally checked the response to the VEGF stimulation for 3 hrs., i.e., the approximate duration of a single peak of Src oscillation. The model predicted and experiments confirmed that this transient VEGF input indeed resulted in a single peak of Src phosphorylation and in a complete recovery of both TEM4 and YAP activity within 4 hrs of VEGF washout. This was in sharp contrast to a much more persistent changes in TEM4 and YAP in response to 2 or 3 peaks of Src phosphorylation triggered by a more prolonged VEGF stimulus (Figs. 4I and S5F). This result suggested that a transient VEGF stimulus can propagate downstream in the signaling pathway, but it leads to a faster adaptation throughout the signaling pathway layers vs. the slower adaptation to more persistent stimuli. The transient nature of each Src peak can thus represent the ability for the pathway to adapt to a transient stimulus, limiting inappropriate pathway stimulation.

### YAP activation promotes upregulation of Dll4 and Jag1, with Jag1 dynamically inhibited due to Dll4 expression

The analysis so far has suggested that two distinct phenotypes linked to VEGF inputs: a) the periodic monolayer re-organization through increased relative cell repositioning and b) the endothelial barrier function, are dynamically controlled on different time scales, defined by Src and TEM4 activation kinetics respectively. Furthermore, the TEM4 dynamics is defined by both a faster-scale Src inputs and a slower-scale YAP feedback. We were interested to see if there may be another layer of regulation downstream of TEM4 and YAP, related to specification of Tip and Stalk cell fates, which is a key pre-requisite for VEGF-induced angiogenesis. Notch signaling is conserved across multicellular eukaryotic organisms, as one of the determinants for cell fate specification. Ligand binding induces cleavage of Notch receptors and release the Notch intracellular domain (NICD) which functions as a transcriptional regulator (26). VEGFR2 signaling promotes Tip cells by inducing the expression of a key Notch ligand, Dll4 (27). Dll4 mediated Notch activation marks the Stalk cells and upregulates VEGFR1 expression acting as a decoy receptor for VEGFA, consequently inhibiting Tip cell fate by suppressing VEGFR2 signaling. Jag1 is another potent Notch ligand crucial for angiogenesis, but it can antagonize Dll4 effects, since it can compete with Dll4 for the receptor binding and has relatively lower signaling capacity (28).

To evaluate potential effect of YAP activation on Dll4 and Jag1, we assayed YAP, Dll4 and Jag1 expression levels in the same cell population. Dynamic analysis of pYAP and the Notch ligands, following VEGF treatment revealed a negative correlation between pYAP and both Dll4 and Jag1 in the first 5 hours of VEGF input (Fig. 5A, S6A,B), suggesting that YAP activity may be involved in the control of expression of both Notch ligands. To further explore this, we repeated the experiment in the presence of 1 μg/ml Verteporfin. We found that the abundance levels of both Dll4 and Jag1 were significantly reduced following YAP inhibition, again supporting a key role of YAP in the control of these Notch ligands (Fig. 5B). We then contrasted the results of the acute YAP inhibition with a long-term genetic perturbation using YAP shRNA (HUVECs and HEK293T cells) and YAP overexpression in HUVECs. The results suggested that both protein and mRNA expression level of Dll4 are down-regulated in cells in response to silencing of YAP expression and, conversely, overexpression of YAP-Flag led to up-regulation of Dll4, indicating that YAP is involved in the transcriptional control of Dll4 (Fig. S6C,D). However, interestingly, the outcome was opposite for Jag1, with evidence pointing to its long-term transcriptional suppression, either direct or indirect, in response to YAP up-regulation.

**Figure 5.**
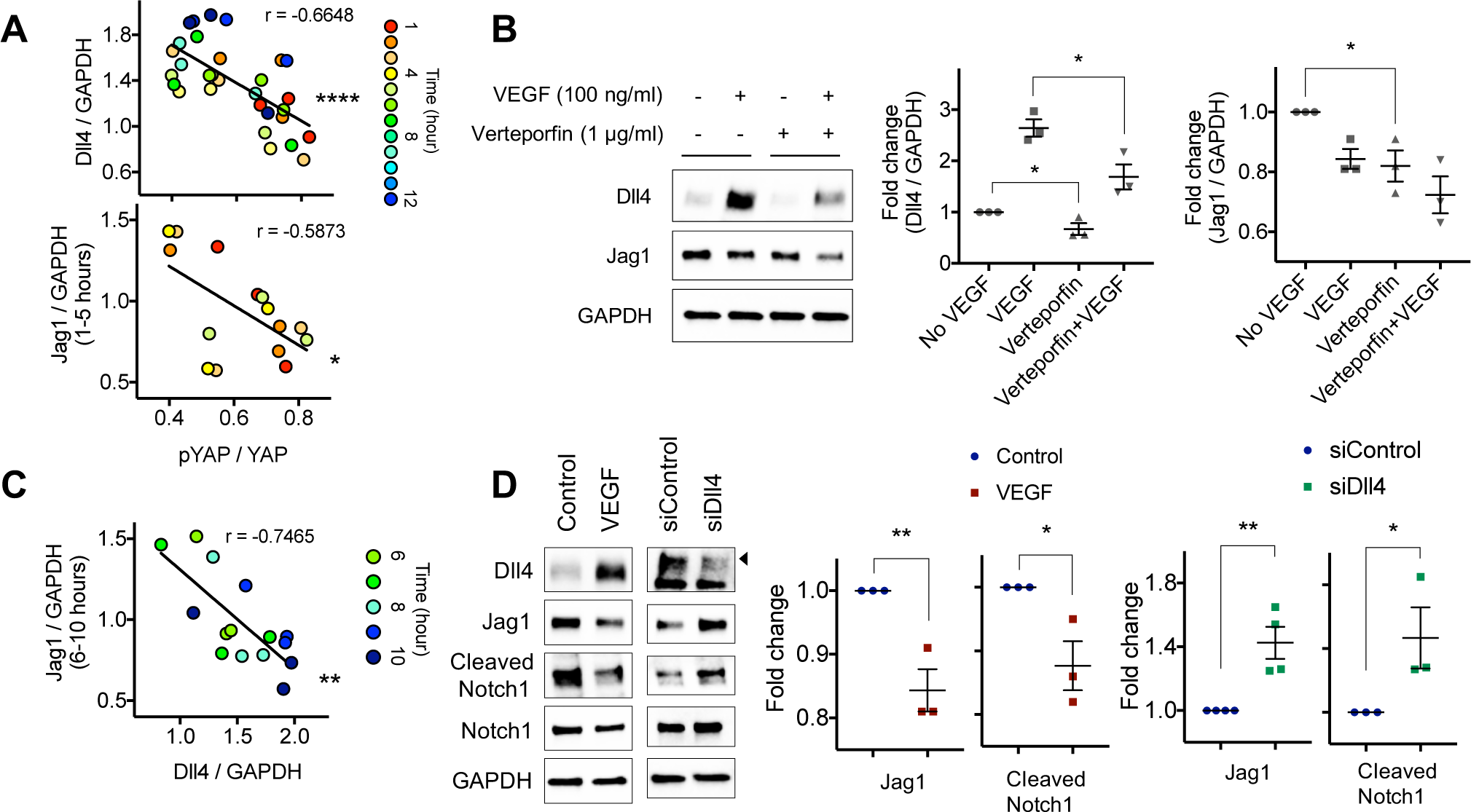
YAP transcriptional activity is involved in the control of the Notch ligands, while Dll4 level inhibiting Jag1 level. **(A)** Scatter plot of fold changes of Dll4/GAPDH **(top)** and Jag1/GAPDH **(bottom)** over pYAP/YAP, based on immuno-blot results including **Figure S4D** and **Figure S6A** (****P<0.0001, *P=0.0213). Color indicates time exposed to VEGF 100 ng/ml. **(B)** Representative immuno-blot results of Dll4 and Jag1 in response to the 1-hour pretreatment of DMSO or Verteporfin 1 μg/ml, followed by the treatment of VEGF 100 ng/ml for 24 hours, and fold change analysis. GAPDH was used as a loading control. (For Dll4/GAPDH, No VEGF vs. Verteporfin only, *P=0.0455, VEGF vs. Verteporfin + VEGF, *P=0.0325, for Jag1/GAPDH, *P=0.0257, n=3) **(C)** Scatter plot of fold changes of Jag1/GAPDH over Dll4/GAPDH, based on immuno-blot results including **Figure S6A** (**P=0.0014). Color indicates time exposed to VEGF 100 ng/ml. **(D)** Representative immuno-blot results of Dll4, Jag1, Cleaved Notch1 and Notch1 of HUVECs treated with VEGF 100 ng/ml for 24 hours, and transfected with Control siRNA, siDll4, and fold change analysis. GAPDH was used as a loading control (For Control vs. VEGF, Jag1/GAPDH, **P=0.0093, n=4, Cleaved Notch1/Notch1, *P=0.0326, n=3, and for Control siRNA vs. siDll4, Jag1/GAPDH, **P=0.0093, n=3, Cleaved Notch1/Notch1, *P=0.0326, n=3)

Our results paradoxically suggested that Jag1 expression is positively controlled by YAP up-regulation on a shorter time scale, but also — negatively over a long-term VEGF stimulation, and thus raised the question of possible underlying mechanisms. The dynamics of Dll4 and Jag1 expression in response to VEGF showed that, although there was a concurrent rise in the abundances of both proteins over the first 6 hrs of VEGF stimulation, Jag1 started to decrease in the 6-12 hrs period., whereas Dll4 continued increasing (Fig. 5C, S6A,B). After 24 hrs. of VEGF stimulation, this divergence increased further, with relatively higher levels of Dll4, being accompanied by lower Jag1 expression, but also lower activated Notch (NICD), suggesting that Dll4 inhibited rather activated Notch signaling at this high levels (Fig. 5D). This inhibitory effect is consistent with the reports that over-expression of Dll4 can inhibit angiogenic sprouting (29). These observations raised the possibility that Dll4, if it exceeds a certain level, can have a negative effect on both Jag1 expression and Notch-mediated cell fate specification. This hypothesis was supported by a relative increase in Jag1 protein expression and NCID abundance in cells expressing siRNA targeting Dll4 (Fig. 5D).

### Signaling frequencies within the VEGF-Src-activated signaling network are filtered out, regulating the expression of Notch ligands

Given the complexity of the VEGF-Src-activated signaling network that we have characterized so far, we further extended the computational model to include the regulation of Dll4 and Jag1 downstream of YAP (Fig. 6A,C, S8A). The extended model was trained on the data from the first 12 hrs. of VEGF stimulation showing excellent agreement with the experimental results. We then validated the model by making further predictions for experimental data not used in the model training. First, we made predictions for the dynamics of signaling of the network components over 60 hrs of VEGF stimulation. The model predicted that over this time, there will be two cycles of a damped TEM4 and YAP oscillations, accompanied by a mostly steady increase in Dll4 and decrease in Jag1 abundances (Fig. 6B). These predictions closely agreed with the experimental analysis over 60 hrs. of VEGF stimulation, providing further support for the model and the underlying mechanism. Furthermore, this result more explicitly revealed two oscillatory processes that occur on different time scales, the first — at the level of Src activation (3 hr. period) and the second — on the level of YAP-TEM4 (24-30 hrs. period), with Dll4 and Jag1 abundances integrated with an even slower time scale of Dll4 synthesis (longer than 60 hrs. time scale). Since Src activated the YAP-TEM4 feedback loop, the faster frequency oscillations (reflecting the Src kinetics) were mixed with slower frequency ones, as reflected by the visible high frequency fluctuations and the Fourier analysis (Fig. 6B, S7A). However, as the signal propagates downstream in the pathway, the slower frequency becomes dominant.

**Figure 6.**
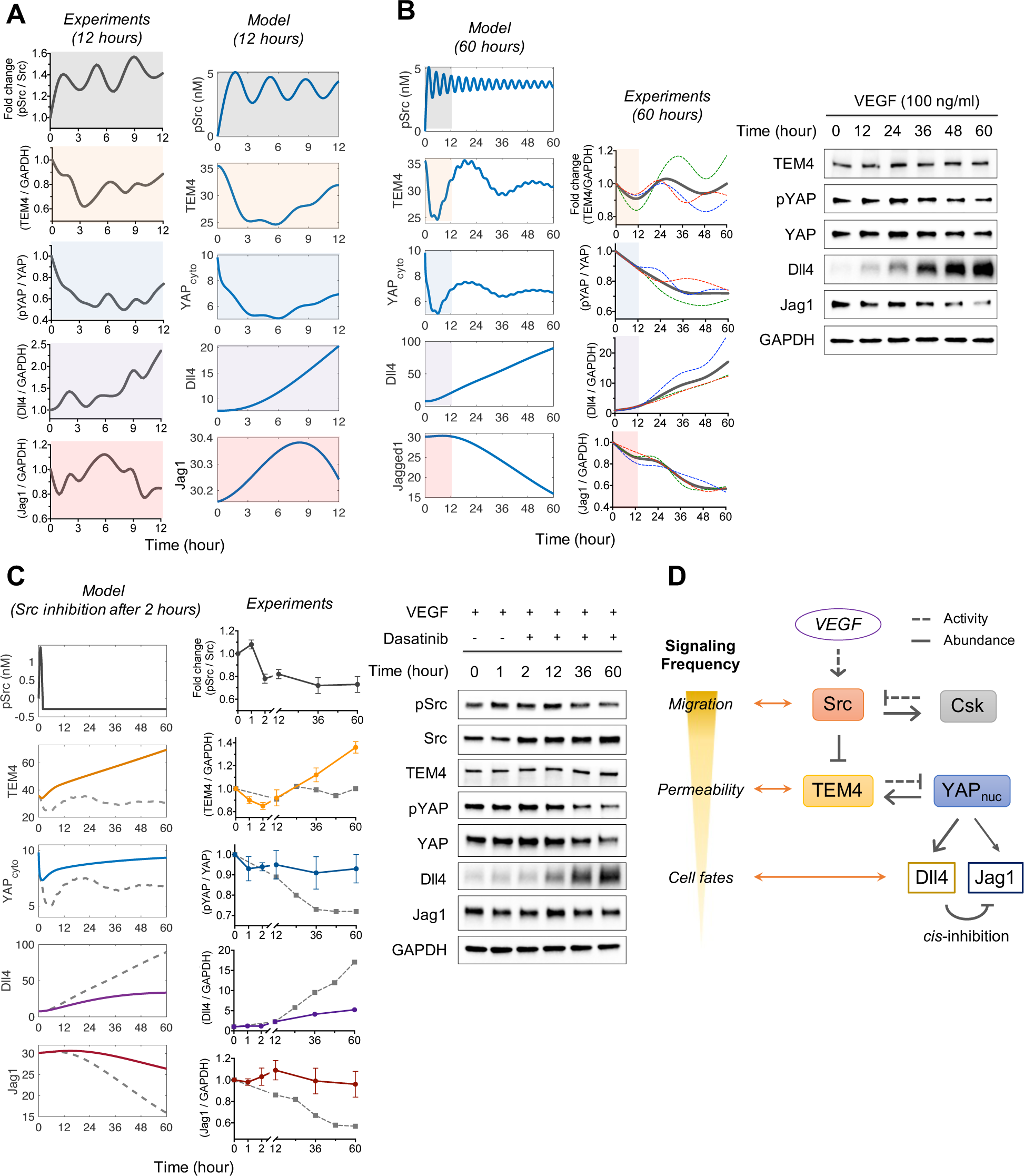
A single input, VEGF, regulates multiple biological outputs by tuning frequency of each signaling module. **(A) (left)** Experimental results of pSrc/Src (gray), TEM4/GAPDH (orange), pYAP/YAP (blue), Dll4/GAPDH (purple) and Jag1/GAPDH (red) of HUVECs in response to VEGF 100 ng/ml for 12 hours from Figure 1B**, Figure S4D and Figure S6A (right)** Model prediction of pSrc (gray), TEM4 (orange), cytoplasmic YAP (blue), Dll4 (purple) and Jag1 (red) for 12 hours when VEGF 2.5 μM (equivalent to 100 ng/ml) is present. **(B) (left)** Model prediction for 60 hours in the same condition, **(middle)** Fold change analysis. Background color corresponds to the 12 hour time portion. Each color dotted line represents individual experiments and gray lines indicate the average. (n=3) **(right)** Representative immuno-blot results of TEM4, pYAP (Ser127), YAP, Dll4 and Jag1. GAPDH was used as a loading control. **(C) (left)** Model prediction of pSrc (gray), TEM4 (yellow), cytoplasmic YAP (blue), Dll4 (purple) and Jag1 (red) for 60 hours in the presence of VEGF 2.5 μM. pSrc is inhibited at 2 hours. **(middle)** Fold change of pSrc/Src, TEM4/GAPDH, pYAP/YAP, Dll4/GAPDH and Jag1/GAPDH of HUVECs for 60 hours in the treatment of Dasatinib 10 nM at 1 hour, in response to VEGF 100 ng/ml. Dotted gray lines indicate the VEGF condition without treatment. (n=4) **(right)** Representative immuno-blot results of TEM4, pYAP (Ser127), YAP, Dll4 and Jag1. GAPDH was used as a loading control. **(D)** Schematic diagram summarizing VEGF induced signaling network controlling multiple phenotypes by the Src-TEM4-YAP pathway.

To further understand the effects of pSrc on the signaling network, we next used the model to predict how a dynamic inactivation of the key upstream input — the activation of Src — would influence the dynamics of signal propagation. In particular, we modeled how inhibition of Src following 1 hr. of VEGF input would affect the signaling outcomes at different pathway steps in the continuous long-term presence of both VEGF and the Src inhibitor. The model predicted that the key effects of this stimulation would be a very transient and low-amplitude activation of the pathway, followed by a progressive increase in TEM4 abundance due to an inhibited Src input, which in turn leads to a transient, low-level YAP activation (Fig. 6D). Surprisingly, even this low-level input led to a considerable (5-fold) increase in Dll4 abundance at 60 hrs. following the initial stimulation. This increase was much lower than the one seen in the absence of Src inhibitor (greater than 15-fold), but was still quite substantial and indicative of the long-term effect of even relatively low level pathway activation due to slow down-regulation and thus long-term ‘memory’ of Dll4 expression, at a far downstream pathway step. The experimental analysis fully supported this prediction, providing yet another validation for the computational model and the assumed underlying regulatory mechanisms. Furthermore, more specially, this agreement between the model and the experiment, supported the assumption that the effects of VEGF signaling on downstream pathway members occur primarily in the Src-dependent fashion rather than through alternative VEGF-dependent pathways.

### Perturbation of the signaling network affects the sprouting angiogenesis

Our results suggest a key role of TEM4 as a downstream Src target in regulation of not only the barrier function (permeability) of the endothelial monolayer, but also, through its regulation of YAP and Notch ligands, of angiogenesis. We tested this putative novel TEM4 function in angiogenesis using a recently proposed validated 3D angiogenesis modeling platform (30), and taking advantage of the experimental results presented above. First we took advantage of the observation that TEM4 expression is elevated if Src is inhibited in the presence of VEGF (Fig. 6C). When we analyzed angiogenesis in the presence of VEGF and dasatinib, we indeed observed reduction in the number and length of sprouts (Fig. S9A,B), indicating a negative effect of TEM4 stabilization. We then directly perturbed TEM4 through shRNA-mediated knockdown and contrasted the angiogenic sprouting in response to VEGF to that exhibited by controlled vessels. Surprisingly, we again observed impaired angiogenesis characterized by shorter sprout length, decreased sprout number, and wider lumens characteristic thicker sprouts (Fig. S9C,D). These results underscored a critical role of TEM4 in angiogenesis and suggested that an optimal level of TEM4 might be needed for effective sprouting, likely due to its dual effect of the control of junctional stability and, through the effect on YAP, angiogenic fate specification.

## Discussion

The analysis integrating modeling and experiments presented in this study reveals a highly dynamic response of Src-dependent signaling in response to VEGF activation. In particular, we find that Src activity is oscillatory, due to a negative feedback control by Csk. This activity directly controls oscillatory changes in the speed of cell migration within the monolayer, leading to relative rearrangement of cells enabling vascular remodeling. Importantly, this activity is further processed by downstream signaling components to control other cellular phenotypes. Importantly, we find that oscillations on Src activity can also translate in oscillatory phosphorylation of VE-cadherin, commonly assumed to control junctional integrity, the permeability of endothelial monolayer is accounted for by a distinct mechanism. Our results suggest that this mechanism relies on the dynamic Src-induced degradation of TEM4, a junctional Rho-GEF protein previously implicated in the maintenance of cellular junctions in both endothelial and epithelial cells (6). Compared to the dynamics of Src phosphorylation cycles, both TEM4 abundance and monolayer permeability (evaluated by measuring the trans-monolayer impedance) show a considerably slower dynamics, determined by the rates of Src-mediated degradation of TEM4 and YAP-mediated TEM4 re-synthesis. Thus we find that two negative feedbacks operate in this Src-activated pathway: one regulating Src activation and involving mutual interaction between Src and Csk, and the other regulating TEM4 abundance, through transcriptional control by YAP. The feedback interaction between TEM4 and YAP is ‘closed’ through the release of YAP from the junctional regions following a decrease in TEM4 and loosening of cellular contacts, allowing YAP to translocate to the cellular nucleus and upregulate TEM4. This YAP activation can also control the expression of other YAP-dependent genes, which, as we show here, include those that encode the fate specifying Notch ligands, Dll4 and Jag1, on an even slower dynamical time scale. Therefore the VEGF-Src-TEM4-YAP pathway described here can indeed regulate distinct cellular phenotypes, including cycles of cell reorganization and cell-contact loosening on a relatively fast time, and gene transcription mediated barrier function and cell fate specification on slower time scales.

The finding of the central role of YAP in regulation of Src-dependent VEGF signaling expands the commonly assumed role of YAP as a sensor of cell-cell contacts and cell density (18, 19) and adds to the growing evidence of involvement of this transcriptional co-regulator in the regulation of vascular function and angiogenesis (31–37). YAP is controlled by angiogenic inducers such as VEGF as well as structural changes in cellular junctions (38). In particular, VEGFR inhibits LATS activity, resulting in activation of YAP (39). Further, loss of function in YAP-TAZ results in irregular endothelial distribution and impaired sprouting formation. Prior observations that YAP is critical for angiogenic responses and barrier function in response to VEGF (33, 35). Our analysis shed further light on functional involvement of YAP in vascular function — as a feedback regulator of junctional contacts, through the control of TEM4 expression, and as a regulator of Dll4 and Jag1 Notch ligands. Furthermore, since YAP responds to mechanical stimuli that can impinge on the endothelium, such as the shear stress from the blood flow (34) and the matrix stiffness (40), it can be a key signaling node integrating multiple input signals to adjust the vascular function and morphogenesis to a variety of micro-environmental conditions.

Signal integration in the VEGF-stimulated network is not limited to YAP activation and likely occurs on multiple other network levels. For instance, our prior results indicate that the VEGF-induced increase in the intracellular calcium levels can directly control oscillatory or persistent activation of MLCK, occurring on a relatively short time scale and regulating cellular motility and cytoskeleton reorganization (41). This activity can be integrated with the control by Src of cell motility within the monolayer, as observed here, as well as contributing to the control of the movement of the Tip cells during the branching morphogenesis. Extensive input integration can also occur at the level of cell fate specification. Transient inputs from the Erk and Akt signaling inputs can be integrated with Src activity in controlling Dll4 expression and promoting the Tip-Stalk cell differentiation. At the same time, we also observe that a longer term input from YAP can lead to a very strong and possibly excessive up-regulation of Dll4 expression that could potentially inhibit Tip cell specification and, as a result, angiogenesis. This result can help reconcile the reports suggesting that Dll4 can be both pro- and anti-angiogenic factor in response to VEGF (28). Indeed, dynamically, there may be a window of time, during which Dll4 levels are sufficiently upregulated to induce effective cell fate specification, but if the input persists, Dll4 would be induced in all, rather than a subset of cells, inhibiting the Tip-Stalk differentiation. The dynamics of cell responses on multiple temporal scales can be a key to a deeper understanding of complex cellular and tissue-level angiogenic responses, integrating multiple signaling inputs.

Our results highlight the role of TEM4 in endothelial function, not only as a regulator of junctional stability, but, through its feedback interaction with YAP, as master regulator of diverse signaling and phenotypic responses downstream of VEGF activated YAP. In particular, our experimental findings in a 3D angiogenesis model system argue that perturbations of TEM4 can lead to profound alterations in formation of vascular sprouts, with both an increase and reduction of the TEM4 expression resulting in impaired angiogenic growth. This findings suggest that an optimal level of TEM4 may be required for coordination of multiple complex processes underlying angiogenesis. In particular, an increase in TEM4 expression may lead to over-stabilization of junctional contacts and inhibition of YAP mediated Dll4 and Jag1 expression, while its decrease may lead a to a decreased stabilization of newly formed and incipient sprouts and, potentially a reduction in cellular contacts required for effective Notch signaling. Further analysis is needed to fully elucidate these mechanisms, but our findings strongly suggest that TEM4 is an interesting target for control of and intervention into vascular function in health and disease.

Our findings raise the question of functional significance of not only the key elements of VEGF activated Src-depend signaling pathway, but also of the oscillatory dynamics controlling their activity. The simulation of the computational model we developed in this study suggest that Src oscillations may not be essential for triggering complex downstream signaling dynamics. Thus we hypothesize that instead, each cycle of the oscillation represents an internal ‘time unit’ representing the continued presence of signaling input. For example, our results suggest that if cells are stimulated transiently, a single cycle of Src activity emerges triggering transient downstream responses, which are in high contrast to much more persistent activation elicited 2 or 3 cycles of Src activity. In other words, Src oscillation may reflect the ability of downstream signaling to quickly reset following a transient and potentially inappropriate inputs, while also being capable of reactivation leading to complex outcomes should the inputs persist over longer time scales.

Overall, our analysis suggests that many processes and phenotypic outputs triggered by VEGF can be encoded in the oscillatory Src activation that is then decoded at different network levels. Deeper levels of the network control longer-term responses, integrating the activity at the preceding upstream levels, whereas the upstream signaling activity is coupled to faster, shorter term phenotypic outputs. The presence of auto-regulatory feedback interactions at each of the network levels can both limit the activity following the input cessation and lead to oscillatory responses on different time scales. In a systems analysis and signal processing sense, one can say that the ‘bandwidth ’ of the signal becomes progressively narrower as the signal propagates through additional steps of the network, with the higher frequencies of the signal being progressively filtered out. This may help not only extend the integration time window (due to a slower signaling response) but also help reduce the effect of stochastic dynamic fluctuations (or signaling noise) that may accumulate in activation of multi-step signaling pathways (42). These findings raise the question of whether similar diverse dynamic response time scales linked to multiple phenotypic outcomes may be found in other signaling pathways and networks.

## Methods and Materials

### Cells and materials

Human umbilical vein endothelial cells (HUVECs) were purchased from Yale Vascular Biology and Therapeutics program and experiments were performed within passage 6. Bovine aortic endothelial cells (BAECs) were a gift of Martin Schwartz’ laboratory at Yale. HEK293T cells were purchased from ATCC (CRL-3216).

### Cell culture

HUVECs were maintained in Medium 199 (Life technologies, 11150-059) with heat inactivated FBS (Life technologies, 16140-071) 20%, Endothelial cell growth supplement 0.03 mg/ml (Sigma-aldrich, E2759), Heparin sodium salts (Sigma-aldrich, H3149-100KU) 0.05 mg/ml, HEPES (Life technologies, 15630106) 10 mM, GlutaMAX supplement (Life technologies, 35050061) 1X and Antibiotic-Antimycotic (Life technologies, 15240112) 1 %. 0.1 % gelatin was coated for 10 minutes at 37 °C incubator before seeding cells on the culture dish. BAECs and HEK293T cells were maintained in DMEM high-glucose media (Corning, 10-013-CV) with heat inactivated FBS 10 % and Penicillin-Streptomycin (Life technologies, 15070063) 1 %. All cell lines were cultured in humidified 37 °C and 5 % CO_2_ incubator.

### Permeability measurement

We used xCELLigence RTCA DP instrument system for non-invasive electrical and real-time impedance measurement. Each glass bottom plate (E-Plate VIEW 16, ACEA Biosciences, 6324738001) was coated with fibronectin 10 μg/ml for 2 hours at room temperature. Cells were seeded and cultured until the impedances reached saturated. Then, cells were serum starved for at least 2 hours before applying VEGF, TNFα or LPA. Serum starved condition for HUVECs is the combination of heat inactivated FBS 1 %, Antibiotic-Antimycotic 1 % and HEPES 10 mM in Medium 199. BAECs were serum starved in DMEM high-glucose media with heat inactivated FBS 1 %, Penicillin-Streptomycin 1 % and HEPES 10 mM. For FITC-dextran assays, we used permeable supports for 12-well plate with 3.0 µm transparent PET (polyethylene terephthalate) membrane (Falcon, 353181). The same procedures were followed for surface coating, cell seeding and serum starving condition with impedance measurement. Fluorescein isothiocyanate-dextran (Sigma, 46945) 0.5 mg/ml was added 1-hour prior to medium collection and fluorescence intensities were analyzed with plate reader (PerkinElmer 2030).

### Immuno-staining and imaging

Cover glass was coated with fibronectin 10 μg/ml for HUVECs and collagen 30 μg/ml for HEK293T cells for 2 hours at room temperature. Cells were washed three times with PBS with MgCl_2_ 1 mM, CaCl_2_ 1 mM and incubated with 4 % paraformaldehyde (PFA) for 15 minutes at room temperature. Fixed cells were permeabilized with 0.1 % Triton X-100 in PBS for 5 minutes and incubated in blocking solution (10 % goat serum in PBS) for 1 hour at room temperature, followed by a treatment with primary antibody overnight at 4 °C. Next day, samples were washed with PBS three times and treated with secondary antibody for 1 hour at room temperature. After washing with PBS three times, samples were applied with ProLong Glass Antifade Mountant (Invitrogen, P36982) and ready for imaging. All imaging experiments were performed with laser scanning confocal microscope (Leica SP8) located in Yale West Campus Imaging Core.

### Immuno-blotting and analysis

Cells were washed three times with PBS with MgCl_2_ 1 mM, CaCl_2_ 1 mM and lysed in RIPA buffer mixed with Halt proteases and phosphatase inhibitor cocktail (Thermo Scientific). Lysed cells were rotated for 10 minutes and centrifuged at 15,000 rpm by table top centrifuge for 10 minutes at 4 °C. After equalizing protein concentration, samples were denatured in 4x Laemmli buffer mixed with 2-mercaptoethanol by boiling at 75 °C for 15 minutes. Subsequently, electrophoresis was performed using Biorad Mini-PROTEAN Tetra system with Mini-PROTEAN precast gels (Biorad) and lysates were transferred onto 0.2 μm nitrocellulose membrane using Tran-Blot Turbo (Biorad) set at 2.5 V, 25 A for 10 minutes. Membranes were incubated in blocking solution (3 % bovine serum albumin in TBST (TBS + 1 % Tween 20)) for 1 hour at room temperature on rotating shaker. Samples were incubated with primary antibody in blocking solution at 4 °C overnight on rotating shaker. Next day, membranes were washed three times with TBST for 15 minutes each and incubated with HRP conjugated secondary antibody in blocking solution for 1 hour at room temperature on rotating shaker. Samples were washed three times again with TBST for 15 minutes each and treated with Biorad Clarity Western ECL substrates to amplify signal. All membrane images were performed in ChemiDoc MP imaging system (Biorad). All immunoblot images were quantified using ImageJ.

### 3D vessel sprouting analysis

Details mimicking sprouting angiogenesis in 3D ECM environment are described elsewhere (30). Briefly, a line mold (diameter of 200 - 250 μm, length of 10 mm) with a semicircular cross-section was deposited on a petri dish using a 3D printer (Ultimaker). A PDMS chamber was put on the mold and a solution of collagen rat tail I (5 mg/ml, BD Biosciences) was injected and allowed to polymerize at 37 °C for 1.5 hours. The mold was then removed and the bottom side of the PDMS chamber was sealed with a glass coverslip. The channel surface was incubated with murine laminin (60 μg/ml, Sigma Aldrich) for 30 minutes at 37 °C with 5 % CO_2_ prior to cell seeding. Suspended HUVECs (5 x 10^5^ cells/ml) were injected through the channel, then incubated at 37 °C with 5 % CO_2_ overnight. Cells were activated with VEGF (100 ng/ml) and TNFα (5 ng/ml) for 3 days. 3D images were acquired with scanning disk confocal microscope (Leica SP8). IMARIS software (Bitplane, Zurich, Switzerland) was used to quantify lumen formation from 3D reconstructed images.

### Co-immunoprecipitation

Co-immunoprecipitation experiment was carried out using Pierce Co-Immunoprecipitation (Co-IP) kit (Thermo Scientific, 26149), according to the manufacturer’s recommendations. Briefly, after swirling the bottle of AminoLink Plus Coupling Resin, 50 μl of the resin slurry was added into a Pierce Spin Column, followed by a centrifugation at 1000 × g for 1 minute. The resin was washed twice with 1X Coupling Buffer, with centrifugation each time. Then, 10 μg of GFP antibody (abcam, ab1218) was prepared to adjust the volume to 200 μl in 1X Coupling Buffer, with addition of 3 μl of the Sodium Cyanoborohydride. The column was incubated on a rotator at room temperature for 2 hours, washed with 1X Coupling Buffer twice. After 200 μl Quenching Buffer was added and centrifuged, the resin was incubated for 15 minutes at room temperature with 200 μl Quenching Buffer + 3 μl of the Sodium Cyanoborohydride. Then, the resin was washed twice by 1X Coupling Buffer, followed by six times washing with 150 μl Wash Solution. After normalizing protein concentration of each lysate in IP Lysis/Wash Buffer, 80 μl of the Control Agarose Resin slurry was added per 1 mg lysate into a spin column and centrifuged, and followed by addition and centrifugation of 100 μl of 1X Coupling Buffer. The lysate was added to the resin and incubated at 4 °C for 1 hour with gentle end-over-end mixing and centrifuged. After the GFP-coupled resin was washed twice with 200 μl IP Lysis/Wash Buffer, the lysate was incubated overnight at 4 °C with gentle rocking. The next day, the sample was washed three times with IP Lysis/Wash Buffer and centrifuged. Then, 10 μl of Elution Buffer was added, centrifuged, and incubated for 5 minutes with 50 μl Elution Buffer at room temperature. The sample was then denatured with the same protocol as immuno-blotting section and proceeded with SDS-PAGE analysis.

### Active Rho GEF pull down assay

Active Rho guanine exchange factor (GEF) pull down assay has been described in detail elsewhere (43). After expressing and purifying GST-RhoA^G17A^ beads, we lysed cells in Rho GEF buffer (20 mM HEPES, 150 mM NaCl, 5 mM MgCl_2_, 1 % (vol/vol) Triton X-100, 1 mM DTT and protease inhibitor cocktail (Sigma)). Then, cells were rotated on shaker for 10 minutes, followed by a centrifugation at 15,000 rpm for 10 minutes at 4 °C. After equalizing protein concentration, 10 μl of GST-RhoA^G17A^ beads were added in each lysates and samples were end-to-end rotated for 45 minutes at 4 °C. After washing the beads three times with Rho GEF buffer, the supernatants were carefully aspirated and 10 μl of Rho GEF buffer was added. Then, samples were denatured in 2x Laemmli buffer mixed with 2-mercaptoethanol by boiling at 75 °C for 15 minutes and SDS-page analysis was performed as described.

### Plasmid transfection and viral infection

We used QIAprep Spin Miniprep kit (QIAGEN) for purifying plasmids. TurboFect (Thermo Scientific) was used for plasmid transfection and Lipofectamine RNAiMAX (Invitrogen) was used for β-catenin siRNA experiment. For viral infection, HEK 293T cells in T75 flask at 70-90 % cell confluency were transfected with psPAX2 4 µg, pMD2.G 2 µg, target vector 4 µg for lentivirus production or MLV gag/pol 4 µg, VSVG 2 µg, target vector 4 µg for retrovirus production (day 0). Cell culture media was refreshed next day and supernatants were collected twice a day (morning and late afternoon) at day 2 and day 3. Then, the collected supernatants were mixed in PEG solution (final concentration 0.1 g/ml PEG 6000, NaCl 0.3 M in deionized water, autoclaved) and stored at 4 °C overnight more than 12 hours. Then, the supernatant/PEG solution mixture were centrifuged at 1500 × g for 30 minutes at 4 °C and virus were pelleted at the bottom of the tube. After carefully removing supernatants from the tube, 1/10 of pellet volume DMEM with 25 mM HEPES was added and stored at -80 °C. On the day of infection, HUVECs at 70-90 % confluency was treated with fresh culture media with polybrene 10 μg/ml and injected with 5 μl/ml concentrated virus. Antibiotic selection process began two days after virus injection. We used 1 μg/ml puromycin for shRNA lentiviral selection and 400 μg/ml G418 for retroviral selection from HUVECs.

### Real-Time Polymerase Chain Reaction

We used RNAprotect Cell Reagent (QUIGEN) to stabilize nucleic acids in cells. RNeasy Plus Mini kit (QUIGEN) was used for purifying RNA. For reverse transcription reaction, we used iScript Reverse Transcription Supermix (BioRad) and incubated the complete mix with the suggested protocol (priming for 5 mins at 25 °C, reverse transcription for 20 mins at 46 °C and inactivation for 1 min at 95 °C) in a thermal cycler (C1000 Touch, Biorad). For gene expression assays, we used TaqMan Fast Advanced Master Mix (Applied Biosystems) and followed the suggested protocol (incubation for 2 mins at 50 °C, polymerase activation for 20 secs at 95 °C, 40 cycles of PCR denaturing for 5 secs at 95 °C and annealing/extending for 30 secs at 55 °C) in real-time PCR system (CFX384 Touch, Biorad). The results were normalized to GAPDH from the same well. Each experiment was carried out in triplicate from three independent biological replicates.

### Chromatin-immunoprecipitation and analysis

Chromatin-immunoprecipitation experiment was carried out using SimpleChIP Plus Enzymatic Chromatin IP kit (Cell Signaling, 9004), according to the manufacturer’s recommendations. For cell culture cross-linking, 540 μl of 37 % formaldehyde was added to each 15 cm culture dish in 20 ml medium with 80 % confluent HUVECs. The dish was incubated 10 minutes at room temperature, followed by an addition of 2 ml of 10x glycine. Cells were incubated 5 minutes at room temperature, added with 2 ml ice-cold PBS with protease inhibitor (Halt proteases and phosphatase inhibitor cocktail, Thermo Scientific), scraped and stored in 15 ml conical tube. Cell were then centrifuged at 2,000 × g for 5 minutes and supernatants were removed for nuclei preparation and chromatin digestion. Cell were then resuspended in 1 ml ice-cold 1x Buffer A added with dithiothreitol (DTT) and protease inhibitor, incubated on ice for 10 minutes with mixing every 3 minutes. Nuclei was pelleted by centrifugation at 2,000 × g for 5 minutes. The supernatant was removed and the pellet was resuspended in 100 μl 1x Buffer B with DTT, and transferred to 1.5 ml microcentrifuge tube. Then, 0.5 μl of Micrococcal Nuclease was added, and the sample was incubated 20 minutes at 37 °C with frequent mixing to digest DNA to length of approximately 150-900 bp. Digest process was stopped by adding 10 μl of 0.5 M EDTA and placing the tube on ice for 2 minutes. The nuclei were pelleted by centrifugation at 16,000 × g for 1 min and the supernatant was removed. The pellet was resuspended in 100 μl of 1x ChIP Buffer with protease inhibitor and incubated on ice for 10 minutes. The sample was sonicated (QSonica) for 20 seconds (1 second sonication, 1 second pause) at 50 % amplitude and incubated on ice for 30 seconds. This process was repeated for three times to break nuclear membrane completely. Lysates were then centrifuged at 9,400 × g for 10 minutes and the supernatant was transferred into a new tube. For chromatin immuno-precipitation (IP), 400 μl of 1x ChIP Buffer with protease inhibitor was added to the 100 μl of the supernatant per IP reaction. A 2.5 μl sample was transferred to a 1.5 ml tube and stored in -80 °C for 0.5% Input analysis. Then, the antibody with the suggested concentration (IgG (Santa Cruz Biotechnology sc-2027), HistonH3 (Cell Signaling #4620), YAP (Cell Signaling #14074), and TEAD1 (Cell Signaling #12292)) was added per 500 μl sample and incubated overnight at 4 °C with rotation. The next day, 30 μl of ChIP-Grade Protein Agarose Beads was added to each IP reaction and incubated 2 hours at 4 °C with rotation. The beads were centrifuged at 3,400 × g for 1 minute and the supernatant was removed, followed by washing by 1 ml of low salt and incubation at 4 °C for 5 minutes with rotation. The washing step with low salt was repeated twice. Then, 1 ml of high salt wash was added to the beads and the sample was incubated at 4 °C for 5 minutes with rotation, followed by a brief centrifugation at 3,400 × g for 1 minute. For elution, 150 μl of 1x ChIP Elution buffer was added to each sample and the chromatin was eluted from the antibody/protein G agarose beads for 30 minutes at 65 °C with gentle vortexing at 250 rpm. Then, the beads were pelleted by brief a 1-minute centrifugation at 3,400 × g and the supernatant were carefully transferred to a new tube. Then, cross-links were reversed by adding 6 μl 5 M NaCl and 2 μl Proteinase K and incubated 2 hours at 65 °C. For DNA purification, 750 μl DNA Binding Buffer was added to each 150 μl DNA sample and each sample was transferred to a DNA spin column in collection tube, followed by a centrifugation at 18,500 × g for 30 seconds. Finally, 50 μl of DNA Elution Buffer was added to each column and centrifuged at 18,500 × g for 30 seconds to elute DNA. For analysis, standard Polymerase Chain Reaction (PCR) method was used. Per 2 μl DNA sample, the following reaction mixes per 1 PCR reaction (18 μl) was used: Nuclease free water 13.1 μl, 10x PCR Buffer 2 μl, 2.5 mM dNTP mix 1.6 μl, 5 μM RPL30 Primer (Cell Signaling #7014) 2 μl or 100 μM target primers 0.4 μl and Taq DNA Polymerase 0.5 μl with the following PCR reaction program (Initial denaturation 95 °C for 5 minutes, denaturation 95 °C for 30 seconds, annealing 62 °C for 30 seconds, extension 72 °C for 30 seconds, with repeat steps for a total of 34 cycles and final extension 72 °C for 5 minutes.) Following sequences of DNA oligo were used for CTGF (forward: 5’-TTGGTGCTGGAAATACTGCG-3’, reverse: 5’-GGGACATTCCTCGCATTCCT-3’) and TEM4 (forward: 5’-CCTGAGTGTCAGTGTGTACGA-3’, reverse: 5’- AGACACACACAACCGCTGAG-3’). Finally, 10 μl of each PCR product was used for analysis by 2 % agarose gel electrophoresis.

### Statistical tests

Data were analyzed by two-tailed t-test, with error bar representing s.e.m, unless otherwise mentioned.

## Acknowledgments

The work was supported by NIH grants U54 CA209992. We thank Andrew Ewald and Alexander Popel for helpful comments.

## Author Contributions

S.H.L. and A.L. conceived the project. S.H.L. designed, performed experiments, and analyzed the data. T.K. performed experiments including TEER measurement, 3D in vitro model, and analyzed the data. S.H.L. and A.L. performed the computational modeling, wrote and edited the manuscript. A.L. obtained the funding, and supervised the project.

