## Supplementary Figures and a Table for "Dynamic decoding of VEGF signaling and coordinated control of multiple phenotypes by the Src-TEM4-YAP pathway"

**Figure S1. Further analysis of VEGF mediated phenotypic processes, related to Figure 1.**
**(A)** Representative immuno-blot results of pSrc (Tyr416), Src, pVE-cadherin (Tyr685), VE-cadherin of HUVECs in response to VEGF 10 ng/ml, and fold change analysis. (n=2 (pSrc/Src). Each color dotted line represents individual experiments and gray lines indicate the average.
**(B)** Representative immuno-blot results of pErk and pAkt in response to VEGF 100 ng/ml, and fold change analysis. GAPDH was used as a loading control. (n=2).
**(C) (top)** Migration speed of HUVECs in response to the 30-min pre-treatment by LY294002 1 μM (blue), PD0325901 1 nM (red) or Dasatinib 10 nM (green), followed by the treatment of VEGF 100 ng/ml. The thin lines represent averaged migration speed from individual cells and the thick lines indicate moving average of the thin lines. **(bottom, left)** Average migration speed for 12 hours of the cells in response to the 30-min pretreatment by LY294002 1 μM, or PD0325901 1 nM, followed by the treatment of VEGF 100 ng/ml. (****P<0.0001, n=12 field of views (FOVs) for all conditions, n=447 cells tracked in average for control (same value from **Figure 1C**, n=433 cells tracked in average for LY294002, n=405 cells tracked in average for PD0325901, all time points included.) **(bottom right)** The number of peaks detected using the peak detection analysis for each condition. (**P=0.0015)
**(D)** Temporal changes of normalized area of HUVECs in response to VEGF 100 ng/ml. Gray lines represent individual measurements. The average is shown in black.
**(E)** Relative real-time impedances of mono-layered BAECs in response VEGF 100 ng/ml. Data were subtracted from control impedance (no treatment, serum-starved) representing baseline 0. Error bar represents s.e.m. (n=3)
**(F)** Representative immuno-blot results of TEM4 in response to VEGF 10 ng/ml, and fold change analysis. (n=3)

**Figure S2. Further analysis for the role of TEM4 at cell-cell junctions, related to Figure 1.**
**(A)** Representative immuno-blot results of TEM4, GEF-H1, p114 RhoGEF, PDZ RhoGEF, Rich1 and p190 RhoGAP in response to VEGF 100 ng/ml for 2 hours, and fold change analysis. GAPDH was used as a loading control. (**P=0.002, n=3)
**(B)** Representative immuno-blot results of co-immunoprecipitation of TEM4 and GEF-H1 against mutant $RhoA^{G17A}$ for mid and high density of HUVECs, and fold change analysis. GAPDH was used as a loading control. (*P=0.016, n=3)
**(C) (top left, bottom)** Representative immuno-blot results, and fold change analysis of TEM4, GEF-H1 and mDia1 for Scrambled shRNA, shGEF-H1 and shTEM4 HUVECs. (***P=0.0001, n=4) **(top, right)** Representative immuno-blot results of TEM4 and mDia1 in HUVECs infected with shTEM4#3 and shTEM4#4. GAPDH was used as a loading control.
**(D)** Representative immuno-staining results, and co-localization analysis of VE-cadherin (green) and F-actin (red) in HUVECs infected with Scrambled shRNA, shGEF-H1 and shTEM4 (scale bar: 50 μm), (**P=0.0027, n=5 (Scrambled shRNA, shGEF-H1) and n=6 (shTEM4))
**(E)** Representative immuno-blot results, and fold change analysis of TEM4, and GEF-H1 over cellular densities. As a loading control, we used β-actin for WCL, EGFR for plasma membrane and GAPDH for cytoplasm. Data were analyzed by two-way ANOVA with error bar representing s.e.m. (****P<0.0001, n=3), **(bottom, right)** Representative immuno-blot results of EGFR, GAPDH and LaminB1 of HUVECs fractionized in nucleus, plasma membrane and cytoplasm.
**(F)** Representative immuno-blot results, and fold change analysis of β-catenin, and TEM4 over increasing doses of β-catenin siRNA from HUVECs. GAPDH was used as a loading control. (*P=0.0179, n=3)
**(G)** Relative real-time impedances of monolayered HUVECs infected with shTEM4 (blue) in response to VEGF 100 ng/ml (red). Data were subtracted from impedance of Scrambled shRNA HUVECs (baseline 0), with error bar representing s.e.m. (n=3).

**Figure S3. Further analysis for the role of pSrc on TEM4 level, related to Figure 3.**
**(A)** Relative real-time impedances of mono-layered HUVECs infected with dominant negative Src mutant, in response VEGF 100 ng/ml. Data were subtracted from control impedance (no treatment, serum-starved) representing baseline with error bar representing s.e.m. (n=3)

**(B)** Representative immuno-blot results of pSrc (Tyr416), Src, pVE-cadherin, VE-cadherin, and TEM4 in response to TNF 10 ng/ml for 4 hours, and fold change analysis. (n=3)

**Figure S4. Further analysis of the role of TEM4 on YAP de-phosphorylation, related to Figure 4.**
**(A)** Representative immuno-staining of YAP (green) and VE-cadherin of HUVECs in response to VEGF 100 ng/ml (scale bar: 100 μm).
**(B)** Representative immuno-blot results of pSrc (Tyr416), Src, pVE-cadherin (Tyr685), VE-cadherin, TEM4, pYAP (Ser127) and YAP of HUVECs in response to VEGF 100 ng/ml for 2 hours, and fold change analysis. GAPDH was used as a loading control. (pSrc/Src, TEM4/GAPDH **P=0.002, pYAP/YAP **P=0.0026, n=3)
**(C)** Representative immuno-blot results of pVE-cadherin (Tyr685), VE-cadherin, TEM4, pYAP (Ser127) and YAP in response to VEGF for 2 hours, and fold change analysis. GAPDH was used as a loading control. Data were analyzed by ordinary one-way ANOVA with error bar representing s.e.m. (For TEM4/GAPDH, *P=0.0447, **P=0.006, ***P=0.0003, for pYAP/YAP, **P=0.0052 (VEGF 10 ng/ml), 0.005 (VEGF 100 ng/ml), n=4)
**(D)** Representative immuno-blot results of pYAP (Ser127) and YAP in response to VEGF 100 ng/ml (left) and 10 ng/ml (right) for 12 hours, and fold change analysis. GAPDH was used as a loading control as in **Figure 1B** and **Figure S1F**. Each color dotted line represents individual experiments and gray lines indicate the average. (n=3)

**Figure S5. Further analysis of the transcriptional role of YAP on the control of TEM4 abundance, related to Figure 4.**
**(A)** Representative immuno-staining of YAP (green) and VE-cadherin of HUVECs in response to LPA 1 μM (scale bar: 100 μm).
**(B)** Representative immuno-blot results of pYAP (Ser127), YAP and TEM4 in response to LPA 1 μM for 2 hours, and fold change analysis. GAPDH was used as a loading control. (pYAP/YAP *P=0.0498, TEM4/GAPDH *P=0.0343, n=3)
**(C)** Representative immuno-blot results of pYAP (Ser127), YAP and TEM4 in response to LPA 1 μM for 2 hours and with pre-treatment of verteporfin 10 μg/ml for 15 hours. GAPDH was used as a loading control. (n=2)
**(D)** Relative real-time impedances of mono-layered HUVECs in response to VEGF 100 ng/ml pre-treated with verteporfin 10 μg/ml for 15 hours. Data were subtracted from impedances of HUVECs pretreated with verteporfin 10 μg/ml (baseline 0) with error bar representing s.e.m. (n=3)
**(E)** Representative immuno-blot results of YAP and TEM4 of HUVECs infected with Scrambled shRNA, shYAP#2, and transfected with Flag-empty, YAP-Flag. GAPDH was used as a loading control. (n=3)
**(F) (left)** Model prediction of temporal kinetics of VEGF (red), pSrc (gray), TEM4 (yellow) and cytoplasmic YAP (blue) when VEGF is removed at 3-hour time point. **(middle)** Fold change analysis of pSrc normalized to Src (gray), TEM4 normalized to GAPDH (yellow), and pYAP normalized to YAP (blue). Gray dotted lines represent the constant VEGF condition. (n=3) **(right)** Representative immuno-blot results of pSrc (Tyr416), Src, pVE-cadherin (Tyr685), VE-cadherin, TEM4, pYAP (Ser127) and YAP of HUVECs in the presence of VEGF 100 ng/ml for the first 3 hours and in the absence of VEGF for the next 4 hours.

**Figure S6. Further analysis of the role of YAP on the control of Dll4 and Jag1, related to Figure 5.**
**(A, B)** Representative immuno-blot results of Dll4 and Jag1 in response to **(A)** VEGF 100 ng/ml (n=3), and **(B)** VEGF 10 ng/ml (n=2) for 12 hours, and fold change analysis. GAPDH was used as a loading control as in **Figure 1B** for (A), and **Figure S1F** for (B). Each color dotted line represents individual experiments and gray lines indicate the average.
**(C)** Representative immuno-blot results of YAP, Dll4 and Jag of HUVECs infected with Scrambled shRNA, shYAP#2, shYAP#1, and of HUVECs transfected with Flag-empty, YAP-Flag and of HEK293T cells infected with Scrambled shRNA and shYAP#2. GAPDH was used as a loading control. (n=3)
**(D)** mRNA relative level of Dll4, Jag1 of HUVECs infected with Scrambled shRNA, shYAP#2, shYAP#1 and of HUVECs transfected with Flag-empty and YAP-Flag. (For Dll4 mRNA level, Scrambled vs. shYAP#2, **P=0.0011, Scrambled vs. shYAP#1, **P=0.0017, for Jag1 mRNA level, **P=0.0012, ****P<0.0001, for Flag-empty vs. YAP-Flag, *P=0.0316, n=3 each)

**(E) (left)** Model prediction of temporal kinetics of Dll4 (purple), and Jag1 (red) when VEGF is removed at 3-hour time point. **(middle)** Fold change of Dll4 normalized to GAPDH (purple), and Jag1 normalized to GAPDH (red). Gray dotted lines represent the constant VEGF condition. (n=3) **(right)** Representative immuno-blot results of Dll4 and Jag1 of HUVECs in the presence of VEGF 100 ng/ml for the first 3 hours and in the absence of VEGF for the next 4 hours.

**Figure S7. Further analysis of VEGF induced temporal kinetics, related to Figure 6.**
**(A)** Fast Fourier Transform analysis based on the time domain results from **Figure 6A,B**. The 12-hour experimental results of pSrc/Src and TEM4/GAPDH from **Figure 1B**. Dotted line represents that the dominant frequency generating the highest amplitude.
**(B)** Representative immuno-blot results of pSrc (Tyr416) and Src of HUVECs for 4 hours in response to VEGF 100 ng/ml with Dasatinib 10 nM treated at 1 hour, and fold change analysis. Error bar represents s.e.m. (n=4)

**Figure S8. Model analysis results, related to Figure 6.**
**(A)** Schematic diagram used in the model, consisting of three modules (1st: red, 2nd: blue, 3rd: green).
**(B)** Sensitivity analysis of the parameters ($k_{1\text{-}5}$) used in the first signaling module.
**(C)** Bifurcation analysis of dominant frequency and its amplitude of pSrc for the parameters ($k_{1\text{-}5}$, $K_{1\text{-}5}$) used in the first signaling module.
**(D)** Pre-equilibrium of TEM4, $YAP_{cyto}$, Dll4 and Jag1 after 60,000 hours simulation in the absence of VEGF input.
**(E)** Bifurcation analysis of dominant frequency and its amplitude of TEM4 for the parameters ($k_{6}$, $k_{8\text{-}10}$, $K_{6}$, $K_{8\text{-}10}$) used in the second signaling module.
**(F)** Bifurcation analysis of the end value of Dll4 and/or Jag1 at 60 hours for the parameters ($k_{11d}$, $k_{12d}$, $k_{11j}$, $k_{12j}$, $k_{13}$ and corresponding *Ks*) used in the third signaling module.
**(G)** Modeling results of various pSrc frequency and the downstream dynamics.
**(H)** Bifurcation analysis of dominant frequency of TEM4, $YAP_{cyto}$, Dll4 and Jag for the parameters, $k_1$ and $K_1$.
**(I)** Bifurcation analysis of averaged value of pSrc, TEM4, $YAP_{cyto}$, Dll4 and Jag1 for the parameters, $k_1$ and $K_1$.
**(J)** Schematic diagram used for pSrc inhibition.

**Figure S9. *In vitro* 3D analysis to study the role of TEM4 in angiogenesis, related to Figure 6.**
**(A)** Representative immuno-staining results of F-actin (green) and nucleus (blue) of HUVECs in response to VEGF 100 ng/ml (control), with the treatment of Dasatinib 10 nM in sprouting vessel. Magnified view from the white dotted boxes and Cross section view from white dotted lines.
**(B) (top)** The number of sprouts and mini-sprouts in each condition. **(bottom)** Length of sprouts or mini-sprouts in each condtion. (**P=, ****P<, Control Sprout n=38, Control Mini-sprout n=100, Dasatinib Sprout n=12, Dasatinib Mini-sprout n=58)
**(C)** Representative immuno-staining results of F-actin (red) and nucleus (blue) of HUVECs infected with Scrambled shRNA or shTEM4 in sprouting vessels (scale bar: 50 $\mu$m). The image intensity in sprouting area is increased to emphasize. (i-ii) Cross section view from white dotted lines. Arrows indicate tip or lumen in sprouting vessels.
**(D) (left, top)** Length of sprouts with lumen for Scrambled shRNA and shTEM4 HUVECs. 1-6 indicates the number of tips per lumen. (**P=0.0018) **(left, middle)** Length of sprouts without lumen for Scrambled shRNA and shTEM4 HUVECs. (**P=0.0052) **(left, bottom)** Density of cells (the number of cells per $\mu$m sprouts) for Scrambled shRNA (n=50) and shTEM4 HUVECs (n=18). (****P<0.0001), **(right)** The number of tip cells and the number of stalk cells from sprouts with or without lumen for Scrambled shRNA and shTEM4 HUVECs.

**Table S1. Model parameters, related to Figure 6.**
**(A)** Summary of velocity of reactions and description used in the model.
**(B)** Summary of equations used in the model.
**(C)** Parameters used in the model.

**Figure S1.**

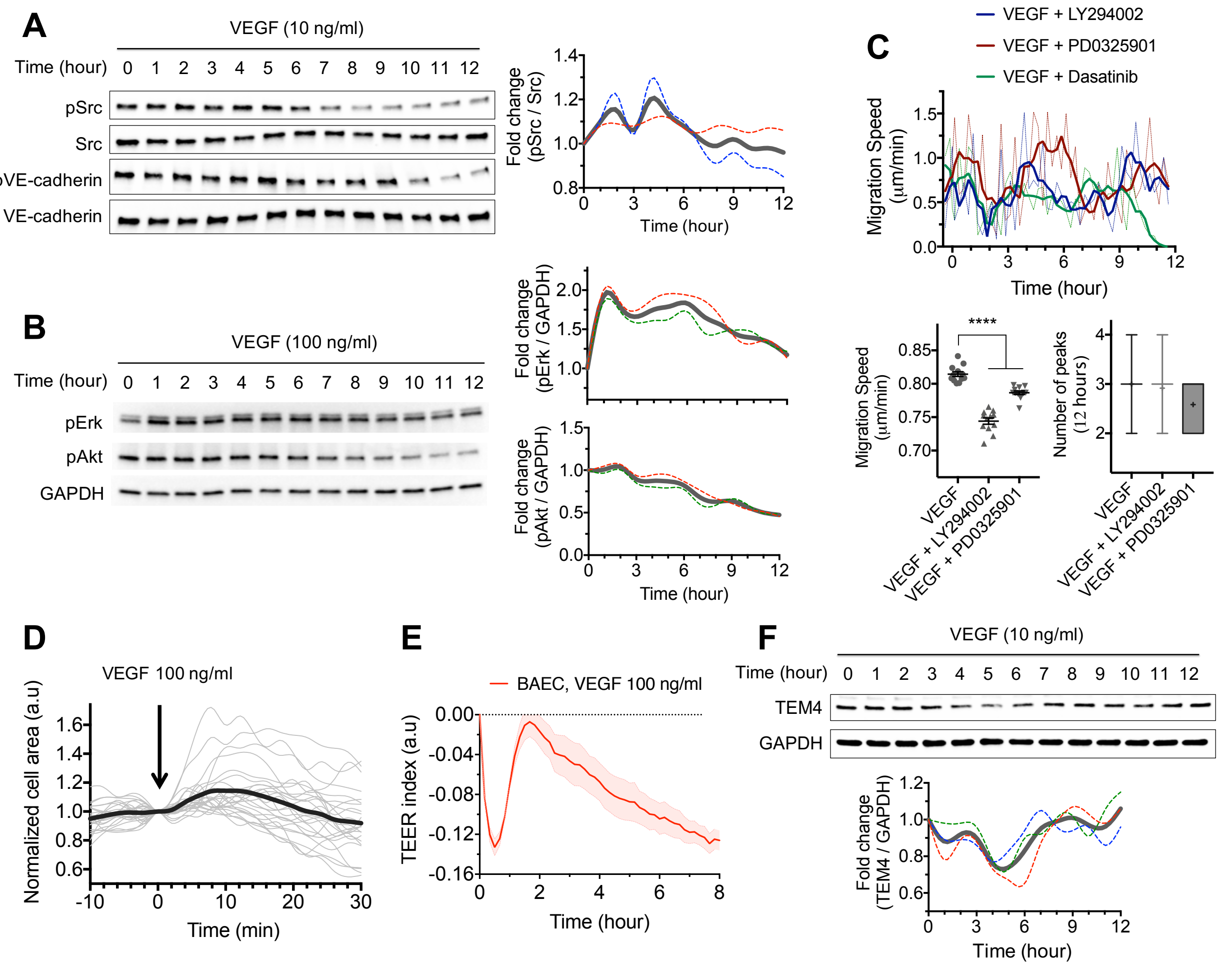
A
VEGF (10 ng/ml)
Time (hour) 0 1 2 3 4 5 6 7 8 9 10 11 12
pSrc
Src
pVE-cadherin
VE-cadherin
Fold change (pSrc / Src)
Time (hour)
B
VEGF (100 ng/ml)
Time (hour) 0 1 2 3 4 5 6 7 8 9 10 11 12
pErk
pAkt
GAPDH
Fold change (pErk / GAPDH)
Fold change (pAkt / GAPDH)
Time (hour)
C
VEGF + LY294002
VEGF + PD0325901
VEGF + Dasatinib
Migration Speed (µm/min)
Time (hour)
****
Number of peaks (12 hours)
VEGF
VEGF + LY294002
VEGF + PD0325901
D
VEGF 100 ng/ml
Normalized cell area (a.u)
Time (min)
E
BAEC, VEGF 100 ng/ml
TEER index (a.u)
Time (hour)
F
VEGF (10 ng/ml)
Time (hour) 0 1 2 3 4 5 6 7 8 9 10 11 12
TEM4
GAPDH
Fold change (TEM4 / GAPDH)
Time (hour)

### Figure S2.

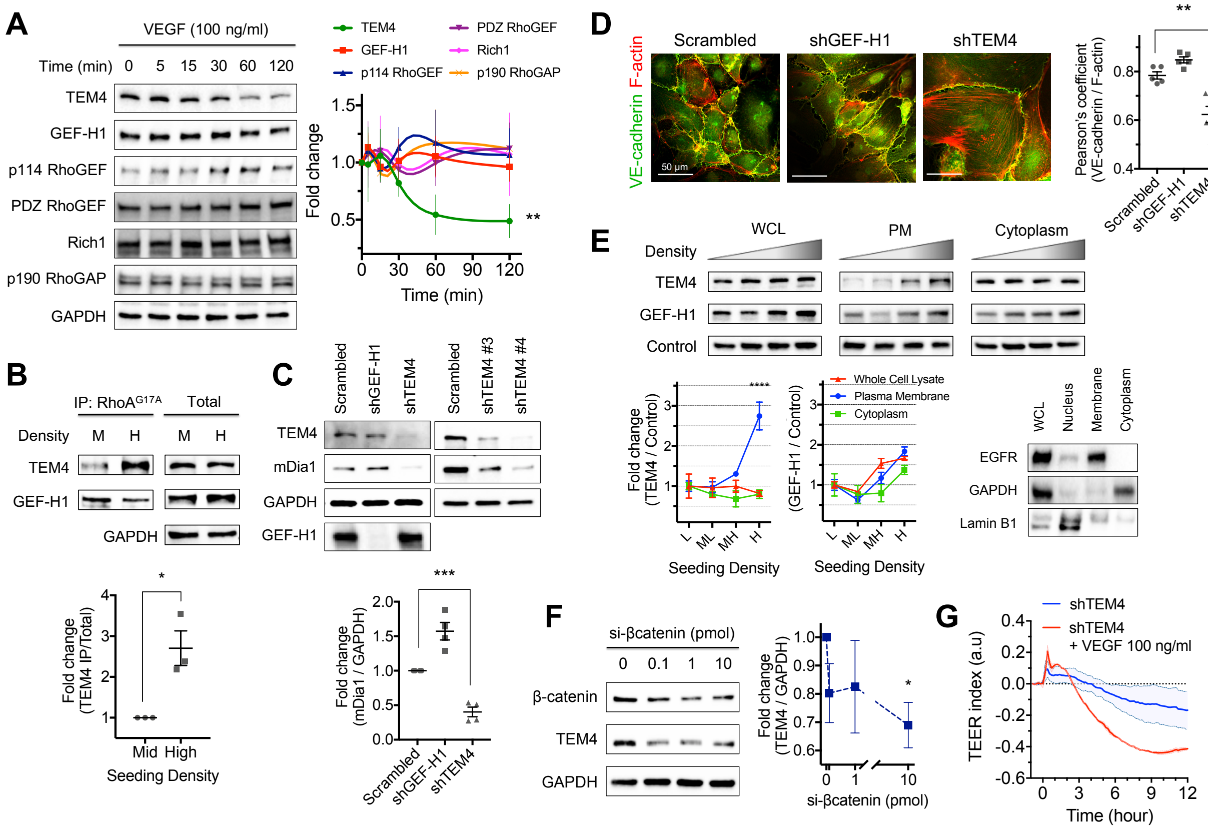


### Figure S3.

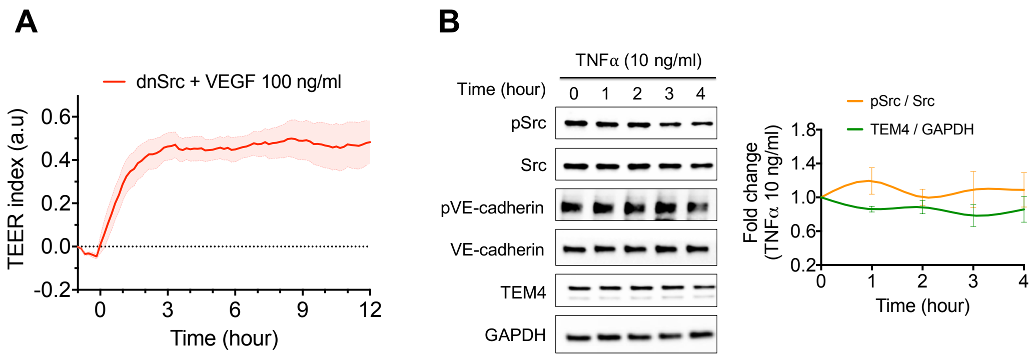

**Figure S4.**

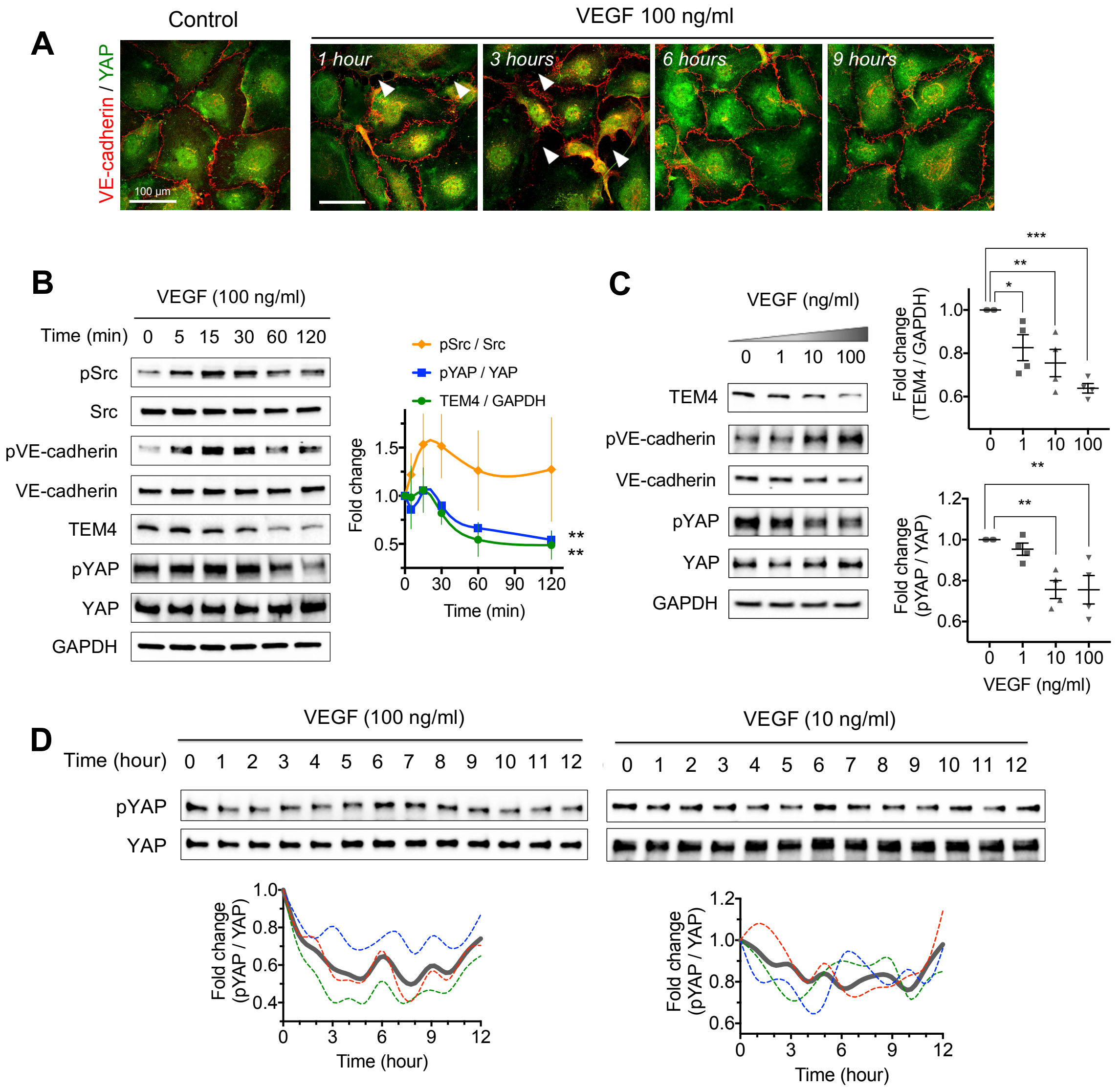

A
Control
VEGF 100 ng/ml
VE-cadherin / YAP
100 µm
1 hour
3 hours
6 hours
9 hours
B
VEGF (100 ng/ml)
Time (min) 0 5 15 30 60 120
pSrc
Src
pVE-cadherin
VE-cadherin
TEM4
pYAP
YAP
GAPDH
pSrc / Src
pYAP / YAP
TEM4 / GAPDH
Fold change
Time (min)
C
VEGF (ng/ml)
0 1 10 100
TEM4
pVE-cadherin
VE-cadherin
pYAP
YAP
GAPDH
Fold change (TEM4 / GAPDH)
Fold change (pYAP / YAP)
VEGF (ng/ml)
D
VEGF (100 ng/ml)
VEGF (10 ng/ml)
Time (hour) 0 1 2 3 4 5 6 7 8 9 10 11 12
pYAP
YAP
Fold change (pYAP / YAP)
Time (hour)

### Figure S5

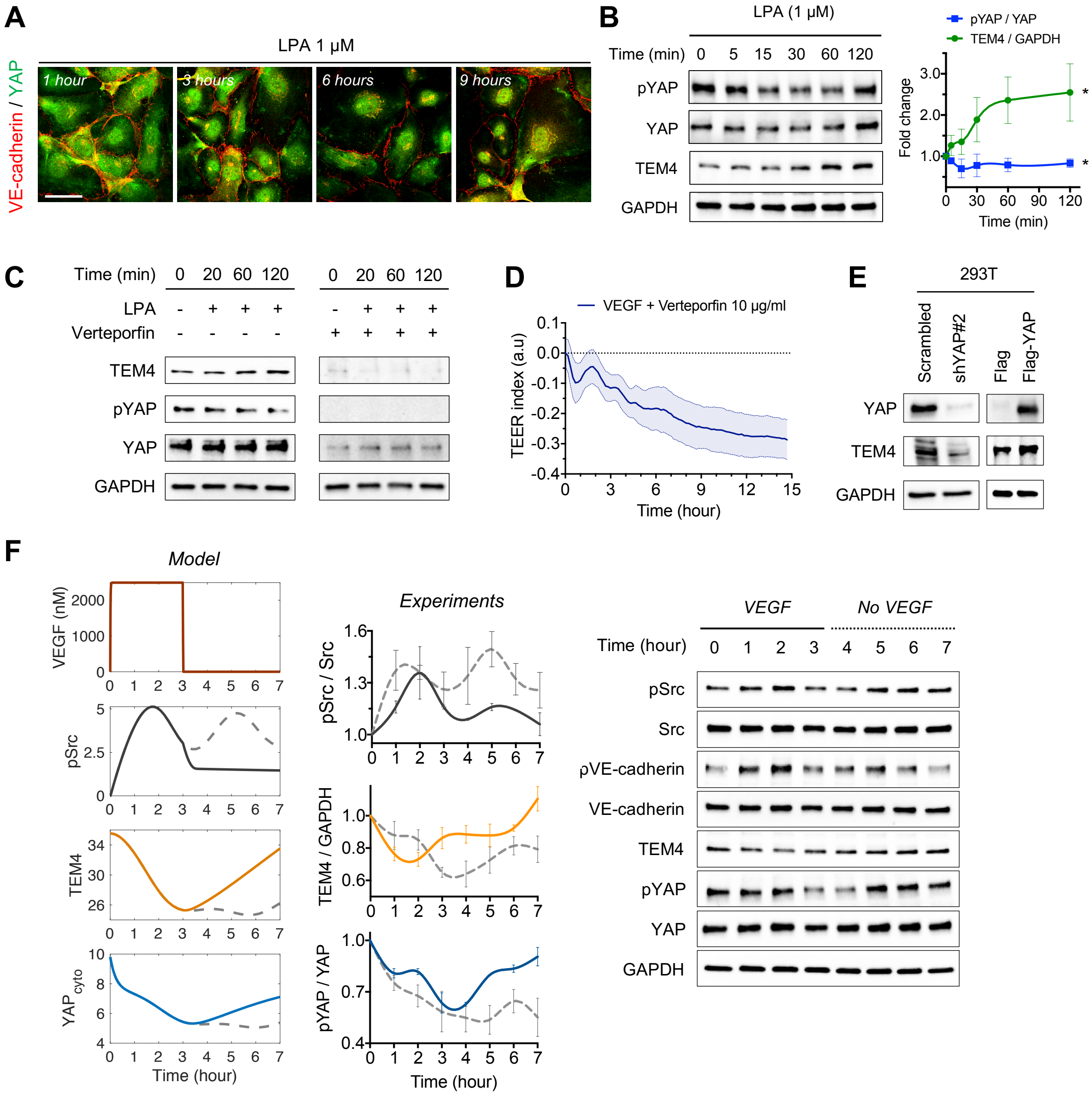

A
LPA 1 µM
VE-cadherin / YAP
1 hour
3 hours
6 hours
9 hours
B
LPA (1 µM)
Time (min) 0 5 15 30 60 120
pYAP
YAP
TEM4
GAPDH
pYAP / YAP
TEM4 / GAPDH
Fold change
Time (min)
C
Time (min) 0 20 60 120 0 20 60 120
LPA
Verteporfin
TEM4
pYAP
YAP
GAPDH
D
VEGF + Verteporfin 10 µg/ml
TEER index (a.u)
Time (hour)
E
293T
Scrambled
shYAP#2
Flag
Flag-YAP
YAP
TEM4
GAPDH
F
Model
VEGF (nM)
pSrc
TEM4
YAPcyto
Time (hour)
Experiments
pSrc / Src
TEM4 / GAPDH
pYAP / YAP
Time (hour)
VEGF
No VEGF
Time (hour) 0 1 2 3 4 5 6 7
pSrc
Src
pVE-cadherin
VE-cadherin
TEM4
pYAP
YAP
GAPDH

**Figure S6.**

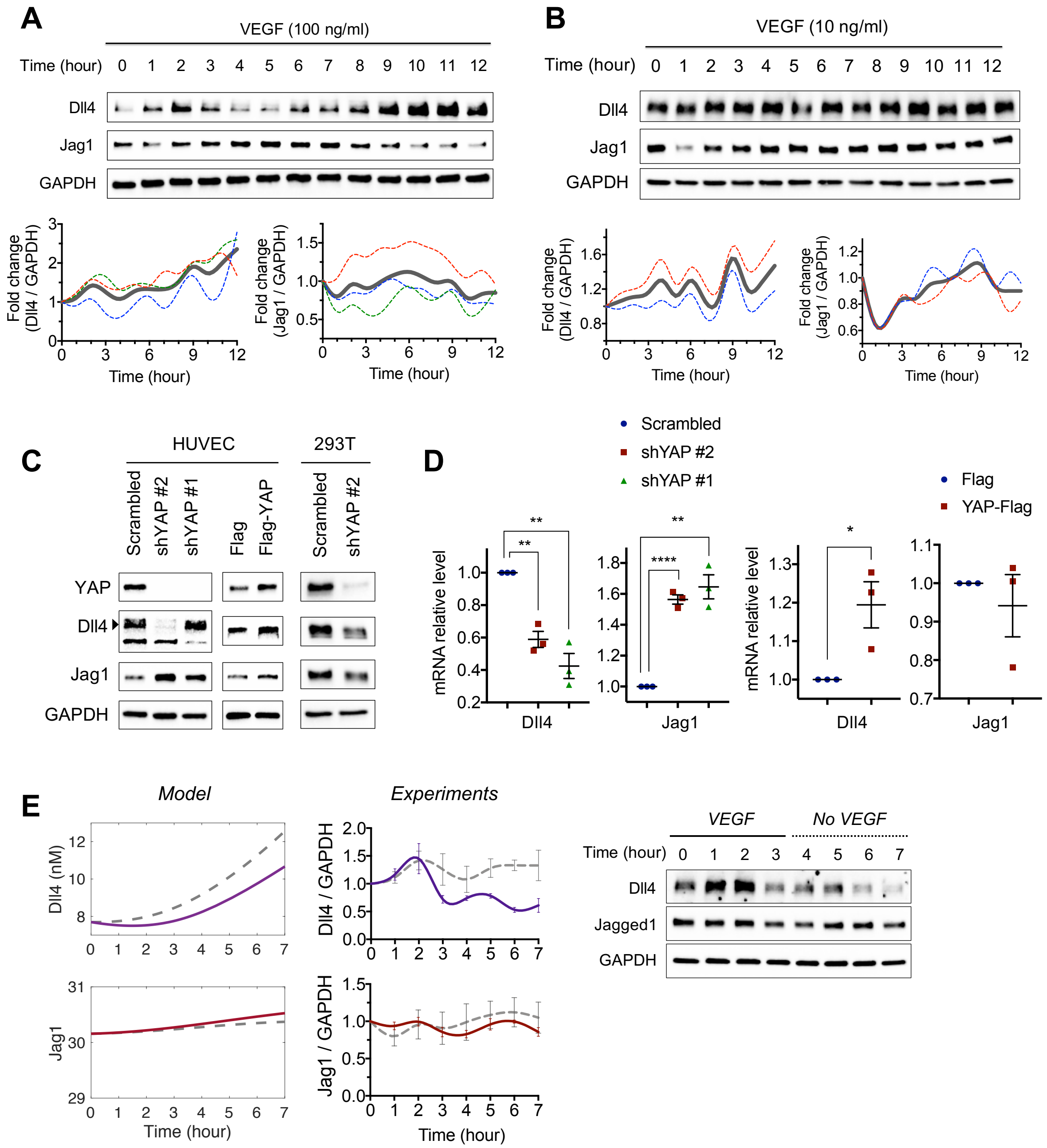

A
VEGF (100 ng/ml)
Time (hour) 0 1 2 3 4 5 6 7 8 9 10 11 12
Dll4
Jag1
GAPDH
Fold change (Dll4 / GAPDH)
Fold change (Jag1 / GAPDH)
Time (hour)
B
VEGF (10 ng/ml)
Time (hour) 0 1 2 3 4 5 6 7 8 9 10 11 12
Dll4
Jag1
GAPDH
Fold change (Dll4 / GAPDH)
Fold change (Jag1 / GAPDH)
Time (hour)
C
HUVEC
293T
Scrambled
shYAP #2
shYAP #1
Flag
Flag-YAP
Scrambled
shYAP #2
YAP
Dll4
Jag1
GAPDH
D
Scrambled
shYAP #2
shYAP #1
Flag
YAP-Flag
mRNA relative level
Dll4
Jag1
E
Model
Experiments
Dll4 (nM)
Jag1
Dll4 / GAPDH
Jag1 / GAPDH
Time (hour)
VEGF
No VEGF
Time (hour) 0 1 2 3 4 5 6 7
Dll4
Jagged1
GAPDH

**Figure S7.**

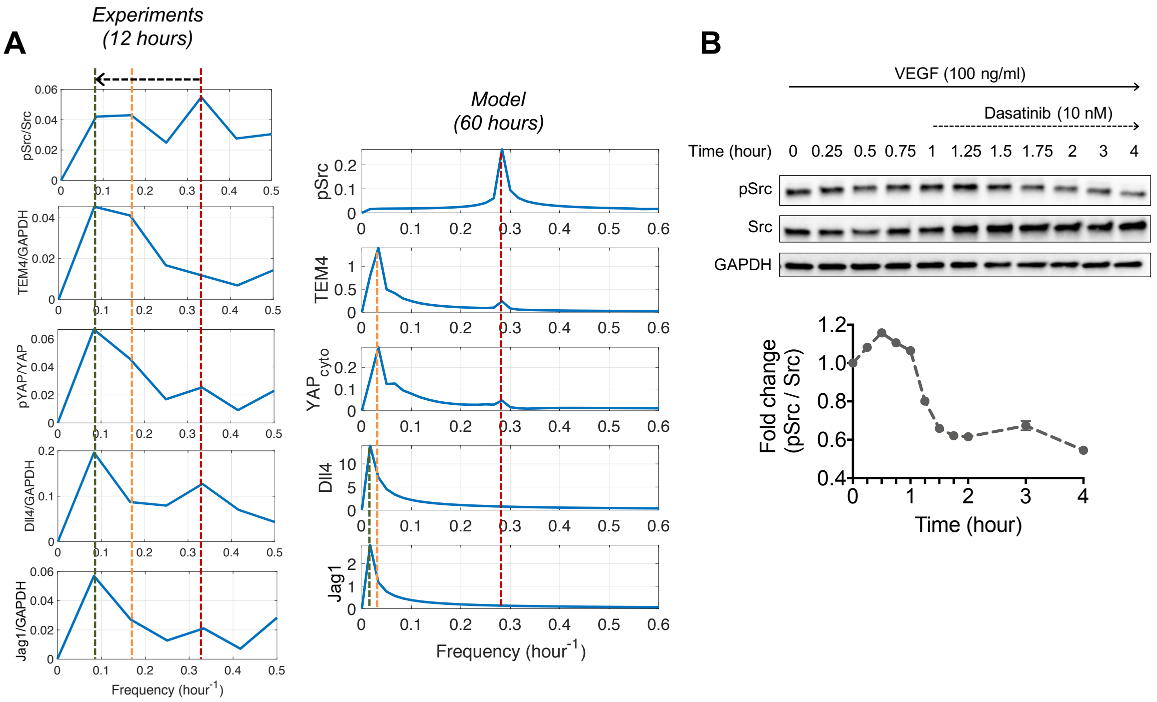
A
Experiments
(12 hours)
pSrc/Src
TEM4/GAPDH
pYAP/YAP
Dll4/GAPDH
Jag1/GAPDH
Frequency (hour$^{-1}$)
Model
(60 hours)
pSrc
TEM4
$YAP_{cyto}$
Dll4
Jag1
Frequency (hour$^{-1}$)
B
VEGF (100 ng/ml)
Dasatinib (10 nM)
Time (hour) 0 0.25 0.5 0.75 1 1.25 1.5 1.75 2 3 4
pSrc
Src
GAPDH
Fold change
(pSrc / Src)
Time (hour)

**Figure S8.**

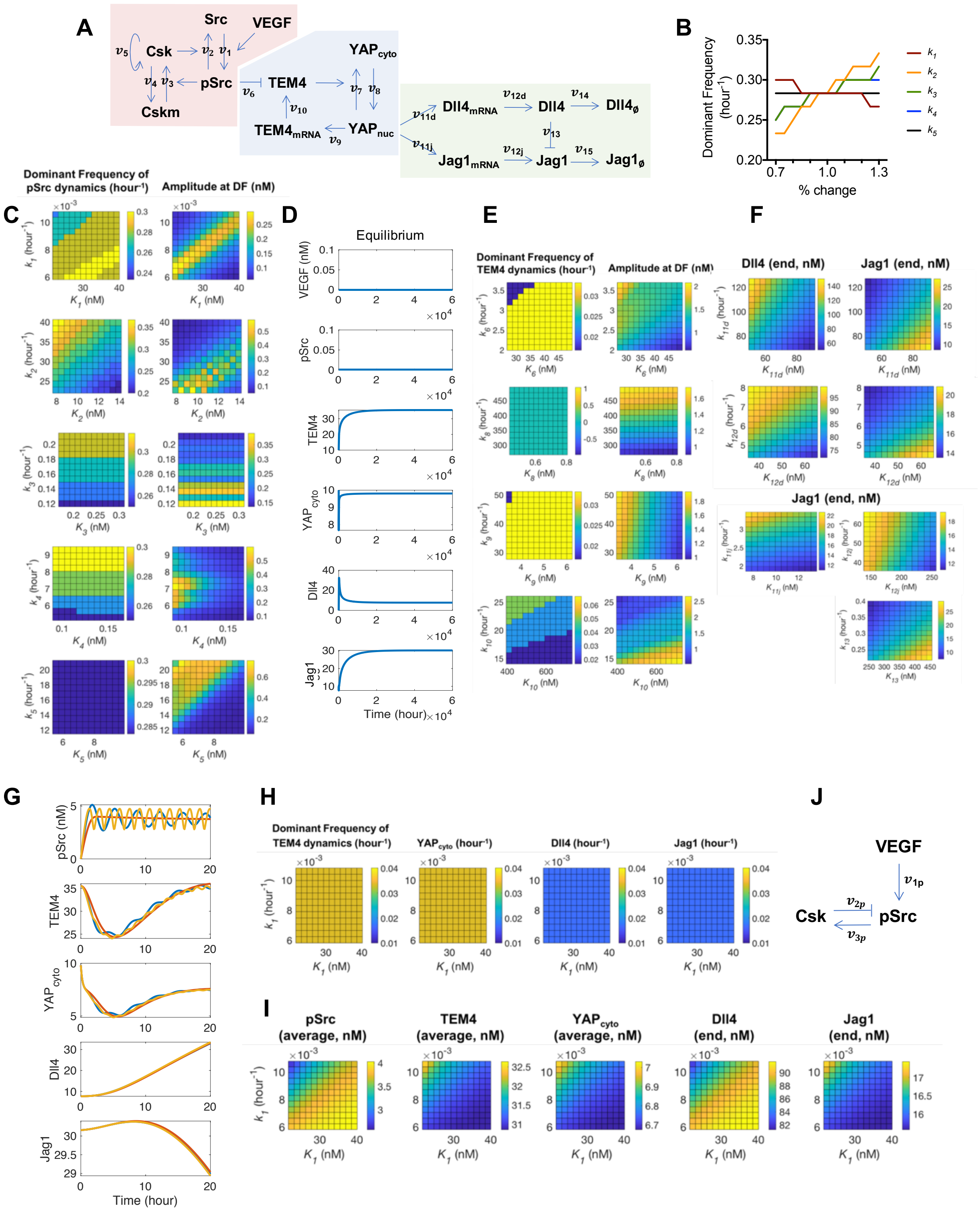
A
Src
VEGF
Csk
pSrc
Cskm
TEM4
YAPcyto
YAPnuc
TEM4mRNA
Dll4mRNA
Dll4
Dll4ø
Jag1mRNA
Jag1
Jag1ø
B
Dominant Frequency (hour-1)
% change
C
Dominant Frequency of pSrc dynamics (hour-1)
Amplitude at DF (nM)
D
Equilibrium
VEGF (nM)
pSrc
TEM4
YAPcyto
Dll4
Jag1
Time (hour)
E
Dominant Frequency of TEM4 dynamics (hour-1)
Amplitude at DF (nM)
F
Dll4 (end, nM)
Jag1 (end, nM)
G
pSrc (nM)
TEM4
YAPcyto
Dll4
Jag1
Time (hour)
H
Dominant Frequency of TEM4 dynamics (hour-1)
YAPcyto (hour-1)
Dll4 (hour-1)
Jag1 (hour-1)
I
pSrc (average, nM)
TEM4 (average, nM)
YAPcyto (average, nM)
Dll4 (end, nM)
Jag1 (end, nM)
J
VEGF
Csk
pSrc

**Figure S9.**

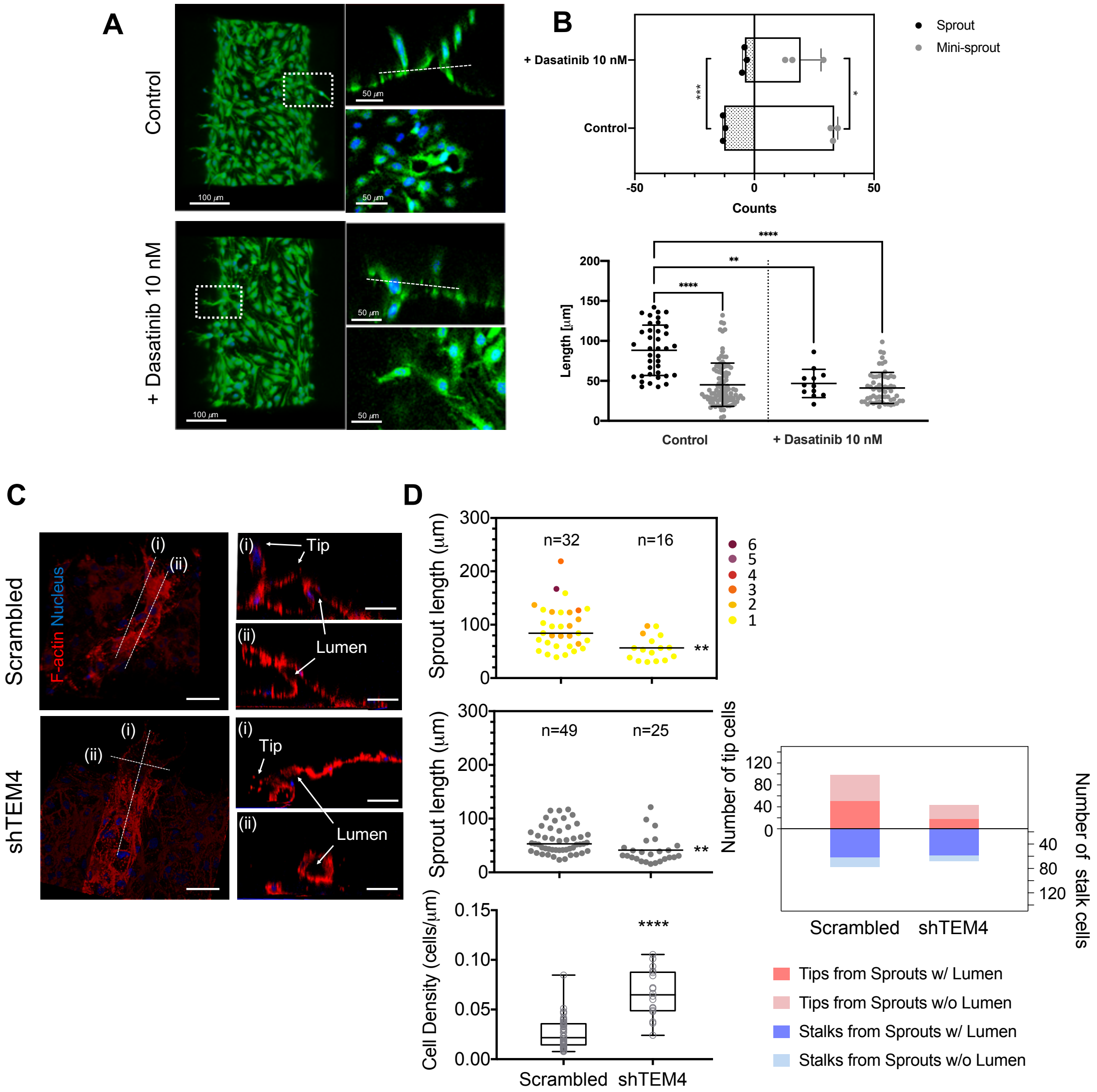

A
Control
+ Dasatinib 10 nM
100 µm
50 µm
B
Sprout
Mini-sprout
+ Dasatinib 10 nM
Control
Counts
Length [µm]
Control
+ Dasatinib 10 nM
C
Scrambled
shTEM4
F-actin Nucleus
Tip
Lumen
D
Sprout length (µm)
n=32
n=16
n=49
n=25
Number of tip cells
Number of stalk cells
Cell Density (cells/µm)
Scrambled
shTEM4
Tips from Sprouts w/ Lumen
Tips from Sprouts w/o Lumen
Stalks from Sprouts w/ Lumen
Stalks from Sprouts w/o Lumen

### Table S1. Model parameters, related to Figure 6.

**A**

| Index | Velocity of reactions | Description |
|---|---|---|
| R1 | $v_1 = \frac{k_1[Src][VEGF]}{K_1+[Src]}$ | the rate of VEGF-dependent Src phosphorylation (pSrc) |
| R2 | $v_2 = \frac{k_2[pSrc][Csk]}{K_2+[pSrc]}$ | the rate of Src dephosphorylation mediated by active Csk |
| R3 | $v_3 = \frac{k_3[Cskm][pSrc]}{K_3+[Cskm]}$ | the rate of Csk activation mediated by pSrc |
| R4 | $v_4 = \frac{k_4[Csk]}{[pSrc](K_4+[Csk])}$ | the rate of pSrc involved Csk inactivation |
| R5 | $v_5 = \frac{k_5[Csk]}{K_5+[Csk]}$ | the rate of Csk activation dependent on itself |
| R6 | $v_6 = \frac{k_6[pSrc][TEM4]}{K_6+[TEM4]}$ | the rate of TEM4 degradation mediated by pSrc |
| R7 | $v_7 = \frac{k_7[TEM4][YAP_{nuc}]}{K_7+[YAP_{nuc}]}$ | the rate of nucleus to cytoplasm trans-localization of YAP mediated by TEM4 abundance |
| R8 | $v_8 = \frac{k_8[YAP_{cyto}]}{[TEM4](K_8+[YAP_{cyto}])}$ | the rate of TEM4 involved cytoplasm to nucleus trans-localization of YAP |
| R9 | $v_9 = \frac{k_9[YAP_{nuc}]^m}{K_9{}^m+[YAP_{nuc}]^m}$ | the rate of *TEM4* transcription mediated by YAP in nucleus ($YAP_{nuc}$) |
| R10 | $v_{10} = \frac{k_{10}[TEM4_{mRNA}]^n}{K_{10}{}^n+[TEM4_{mRNA}]^n}$ | the rate of TEM4 translation |
| R11 | $v_{11d} = \frac{k_{11d}[YAP_{nuc}]^p}{K_{11d}{}^p+[YAP_{nuc}]^p}$ | the rate of *Dll4* transcription mediated by $YAP_{nuc}$ |
| R12 | $v_{12d} = \frac{k_{12d}[Dll4_{mRNA}]^q}{K_{12d}{}^q+[Dll4_{mRNA}]^q}$ | the rate of Dll4 translation |
| R13 | $v_{11j} = \frac{k_{11j}[YAP_{nuc}]^r}{K_{11j}{}^r+[YAP_{nuc}]^r}$ | the rate of *Jagged1* transcription mediated by $YAP_{nuc}$ |
| R14 | $v_{12j} = \frac{k_{12j}[Jag1_{mRNA}]^s}{K_{12j}{}^s+[Jag1_{mRNA}]^s}$ | the rate of Jagged1 translation |
| R15 | $v_{13} = \frac{k_{13}[Jag1][Dll4]}{k_{13}+[Jag1]}$ | the rate of Jagged1 downregulation mediated by Dll4 level |
| R16 | $v_{14} = k_{14}[Dll4]$ | the rate of Dll4 degradation |
| R17 | $v_{15} = k_{15}[Jag1]$ | the rate of Jagged1 degradation |
| R18 | $v_{1p} = \frac{k_1[VEGF]}{K_1+[VEGF]}$ | the rate of VEGF-mediated Src activation |
| R19 | $v_{2p} = \frac{k_2[Csk]}{K_2+[Csk]}$ | the rate of pSrc inhibition mediated by Csk |
| R20 | $v_{3p} = \frac{k_3[pSrc]}{K_3+[pSrc]}$ | the rate of Csk activation mediated by pSrc |

**B**

| Index | Equations |
|---|---|
| E1 | $\frac{d[VEGF]}{dt} = ae^{-\lambda t}$ |
| E2 | $\frac{d[Src]}{dt} = v_2 - v_1$ |
| E3 | $\frac{d[pSrc]}{dt} = v_1 - v_2$ |
| E4 | $\frac{d[Csk]}{dt} = v_3 - v_4 + v_5$ |
| E5 | $\frac{d[Cskm]}{dt} = v_4 - v_3$ |
| E6 | $\frac{d[TEM4]}{dt} = v_{10} - v_6$ |
| E7 | $\frac{d[YAP_{cyto}]}{dt} = v_7 - v_8$ |
| E8 | $\frac{d[YAP_{nuc}]}{dt} = v_8 - v_7$ |
| E9 | $\frac{d[TEM4_{mRNA}]}{dt} = v_9 - v_{10}$ |
| E10 | $\frac{d[Dll4_{mRNA}]}{dt} = v_{11d} - v_{12d}$ |
| E11 | $\frac{d[Dll4]}{dt} = v_{12d} - v_{14}$ |
| E12 | $\frac{d[Jag1_{mRNA}]}{dt} = v_{11j} - v_{12j}$ |
| E13 | $\frac{d[Jag1]}{dt} = v_{12j} - v_{13} - v_{15}$ |
| E14 | $\frac{d[Dll4_{\emptyset}]}{dt} = v_{14}$ |
| E15 | $\frac{d[Jag1_{\emptyset}]}{dt} = v_{15}$ |
| E16 | $\frac{d[VEGF]}{dt} = ae^{-\lambda t}$ |
| E17 | $\frac{d[Csk]}{dt} = v_3$ |
| E18 | $\frac{d[pSrc]}{dt} = v_{1p} - v_{2p}$ $(when\ t \le 2, else\ 0)$ |

**C**

| Parameter | Value (k's: $hr^{-1}$, K's: nM) |
|---|---|
| $a$ | 250000 |
| $\lambda$ | 100 |
| $k_1$ | 0.0083 (Kaimachnikov et al) |
| $K_1$ | 30.8247 |
| $k_2$ | 31.2587 |
| $K_2$ | 10.8939 |
| $k_3$ | 0.1653 |
| $K_3$ | 0.2397 |
| $k_4$ | 7.2945 |
| $K_4$ | 0.1285 |
| $k_5$ | 16.3549 |
| $K_5$ | 7.6367 |
| $k_6$ | 2.8435 (Titsias et al) |
| $K_6$ | 38.3130 |
| $k_7$ | 30*$k_8$ (Sun et al) |
| $K_7$ | 101881.4598 |
| $k_8$ | 381.1336 |
| $K_8$ | 0.6204 |
| $k_9$ | 39.3898 |
| $K_9$ | 4.7921 |
| $m$ | 5.2087 |
| $k_{10}$ | 20.0395 |
| $K_{10}$ | 564.4202 |
| $n$ | 0.4933 |
| $k_{11d}$ | 97.7418 |
| $K_{11d}$ | 70.568 |
| $p$ | 1.2105 |
| $k_{12d}$ | 6.2026 |
| $K_{12d}$ | 50.4093 |
| $q$ | 0.9967 |
| $k_{11j}$ | 2.6613 |
| $K_{11j}$ | 10.4495 |
| $r$ | 0.4645 |
| $k_{12j}$ | 50.834 |
| $K_{12j}$ | 197.45 |
| $s$ | 2.8893 |
| $k_{13}$ | 0.3077 |
| $K_{13}$ | 351.0144 |
| $k_{14}$ | 0.0104 (Titsias et al) |
| $k_{15}$ | 0.0001 (Titsias et al) |
| $k_{1p}$ | 2.85 |
| $K_{1p}$ | 9.59 |
| $k_{2p}$ | 12.72 |
| $K_{2p}$ | 55.2 |
| $k_{3p}$ | 200.07 |
| $K_{3p}$ | 8.5 |
